# Local photo-crosslinking of native tissue matrix regulates cell function

**DOI:** 10.1101/2024.08.10.607417

**Authors:** Donia W. Ahmed, Matthew L. Tan, Jackson Gabbard, Yuchen Liu, Michael M. Hu, Miriam Stevens, Firaol S. Midekssa, Lin Han, Rachel L. Zemans, Brendon M. Baker, Claudia Loebel

**Author notes:** equal contribution (Donia W. Ahmed, Matthew L. Tan).

## Abstract

Within most tissues, the extracellular microenvironment provides mechanical cues that guide cell fate and function. Changes in the extracellular matrix such as aberrant deposition, densification and increased crosslinking are hallmarks of late-stage fibrotic diseases that often lead to organ dysfunction. Biomaterials have been widely used to mimic the mechanical properties of the fibrotic matrix and study cell function. However, the initiation of fibrosis has largely been overlooked, due to the challenges in recapitulating early fibrotic lesions within the native extracellular microenvironment. Using visible light mediated photochemistry, we induced local crosslinking and stiffening of extracellular matrix proteins within *ex vivo* murine and human tissue. In *ex vivo* lung tissue of epithelial cell lineage-traced mice, local matrix crosslinking mimicked early fibrotic lesions that increased alveolar epithelial cell spreading, differentiation and extracellular matrix remodeling. However, inhibition of cytoskeletal tension or integrin engagement reduced epithelial cell spreading and differentiation, resulting in alveolar epithelial cell dedifferentiation and reduced extracellular matrix deposition. Our findings emphasize the role of local extracellular matrix crosslinking and remodeling in early-stage tissue fibrosis and have implications for *ex vivo* disease modeling and applications to other tissues.

## Main

The extracellular matrix (ECM), a complex network of proteins and proteoglycans, provides mechanical signals to constituent cells ^1,2^, and serves as a reservoir of biochemical signaling cues that regulate cell and resultant organ function ^3^. Within the lung, the basement membrane ECM primarily provides support for epithelial and endothelial cells, while the interstitial ECM enables mesenchymal cell adhesion and function central to structural maintenance of the tissue ^4,5^ Changes in ECM structure and mechanical properties are well-described in the developing and diseased lung, including excessive interstitial ECM deposition characteristic of fibrotic scarring ^6,7^. While overall ECM stiffening with loss of lung function is attributed to late-stage fibrosis, localized ECM densification and crosslinking are commonly described as early-stage fibrotic signatures ^8^. These ECM signals are likely important towards mediating epithelial cell fate within the distal lung. For example, increased mechanical strain was recently reported to induce differentiation of cuboidal type 2 alveolar progenitor cells (AT2 cells) into squamous alveolar type 1 cells (AT1 cells) ^9–11^.However, there are no reports specifically examining whether ECM stiffening within the alveolar microenvironment may promote or impair AT2-to-AT1 differentiation.

Within the distal lung epithelium, resident AT2 cells are anchored to the basement membrane where they serve as progenitors to constantly repair alveolar injuries through proliferation and differentiation into AT1 cells ^12,13^. Ineffectual AT2 differentiation, leading to the accumulation of AT2 transitional cells, has now been associated with the formation of fibrotic lesions, suggesting an altered and insufficient repair process ^14–16^. As fibrotic remodeling progresses, ECM is continuously being synthesized, densified, and crosslinked, leading to increased mechanical strain and cytoskeletal tension within alveolar epithelial cells ^17–20^. Thus, dynamic interactions between differentiating AT2 cells and the ECM are a critical component of early-stage fibrotic remodeling. Despite recent evidence suggesting that AT2 transitional cells are characteristic of early fibrotic remodeling, there are no reports on the influence of ECM stiffening on the emergence and persistence of these transitional cells.

Transmission of mechanical changes to the cell is well-established to regulate cell signaling and function, including intracellular contractility, differentiation, and the deposition of newly synthesized ECM ^21–23^. Synthetic and ECM-derived polymeric hydrogels with tunable mechanical properties have been used to recapitulate such mechanical changes *in vitro* ^24–28^. In particular, the use of cyto-compatible photochemistries, such as methacrylate, norbornene and tyrosine-modified polymers, has enabled on demand hydrogel stiffening in the presence of cells towards investigating this critical signal’s potential contribution to changes in cell function ^29,30^. Indeed, dynamic stiffening has been reported to increase cell spreading and intracellular contractility as well as enhance cell differentiation into fibrotic phenotypes such as myofibroblast-like and dedifferentiated epithelial cells of various tissues ^31–33^. However, there are no reports on the role of local ECM stiffening within three-dimensional tissues models to investigate epithelial cell function, suggesting a lack of fundamental knowledge on matrix-driven progression of disease.

To address this, we developed an early-stage *ex vivo* lung fibrosis model using visible light-mediated crosslinking of tyrosine residues to induce local ECM stiffening in murine and human lung tissue slices. This model enables on demand, *in situ* stiffening of the ECM that cells adhere to, allowing us to probe the role of ECM stiffening on induced lung epithelial cell differentiation and function through intracellular and extracellular mechanisms within their native microenvironments. More specifically, our findings show that ECM stiffening increases the deposition of ECM proteins, and that subsequent lung epithelial cell function requires intracellular contractility and integrin engagement.

### Blue light-mediated ECM-derived polymer crosslinking

The alveolar epithelium has an extensive reparative capacity that requires a stem cell driven regenerative response ^34^. In general, following epithelial cell damage, progenitor cell populations, including AT2 cells, are mobilized to promote alveolar regeneration ^35–39^. AT2 cells are capable of both self-renewal and differentiation into AT1 cells to reestablish a functional alveolar epithelium. However, microfoci of repeated cycles of epithelial damage following injury are postulated to contribute to the development of a dysfunctional repair response and subsequent early fibrotic lesions through aberrant ECM deposition, densification and crosslinking which lead to fibrotic remodeling (Figure 1A). To model this, we adopted a ruthenium (Ru)- mediated chemistry towards local crosslinking of tyrosines-containing ECM proteins (Figure 1B). When exposed to blue light, the photosensitizers ruthenium (Ru) and sodium persulfate (SPS) induce Ru(III) and sulfate radicals, initiating the formation of tyrosyl radicals ^40,41^ and subsequent arene coupling with a nearby tyrosine residue or radical isomerization with neighboring tyrosyl radicals^42^ that leads to dityrosine bonds (Supplementary Fig 1). To demonstrate the formation of dityrosine bonds, we used ECM-derived polymer hydrogels with embedded fluorescent beads and particle image velocimetry (PIV) to visualize their displacement upon light exposure of a circular 100 µm diameter region of interest (ROI) (Figure 1C). Commonly used and commercially available ECM-derived polymers including collagen type I, fibrinogen, and gelatin were first polymerized into hydrogels either through temperature-induced self-assembly (collagen type I and gelatin) or enzymatic crosslinking via thrombin (fibrinogen), followed by blue light exposure that induced bead displacement towards the center (Figure 1D). We observed varying degrees of bead displacement due to gel contraction and varying spatial resolution in response to light, presumably because of different initial crosslinking densities and other properties such as porosity of the hydrogels ^43–46^. Dityrosine-induced polymer densification and contractions were also confirmed by brightfield imaging (Supplementary Figure 2).

**Figure 1:**
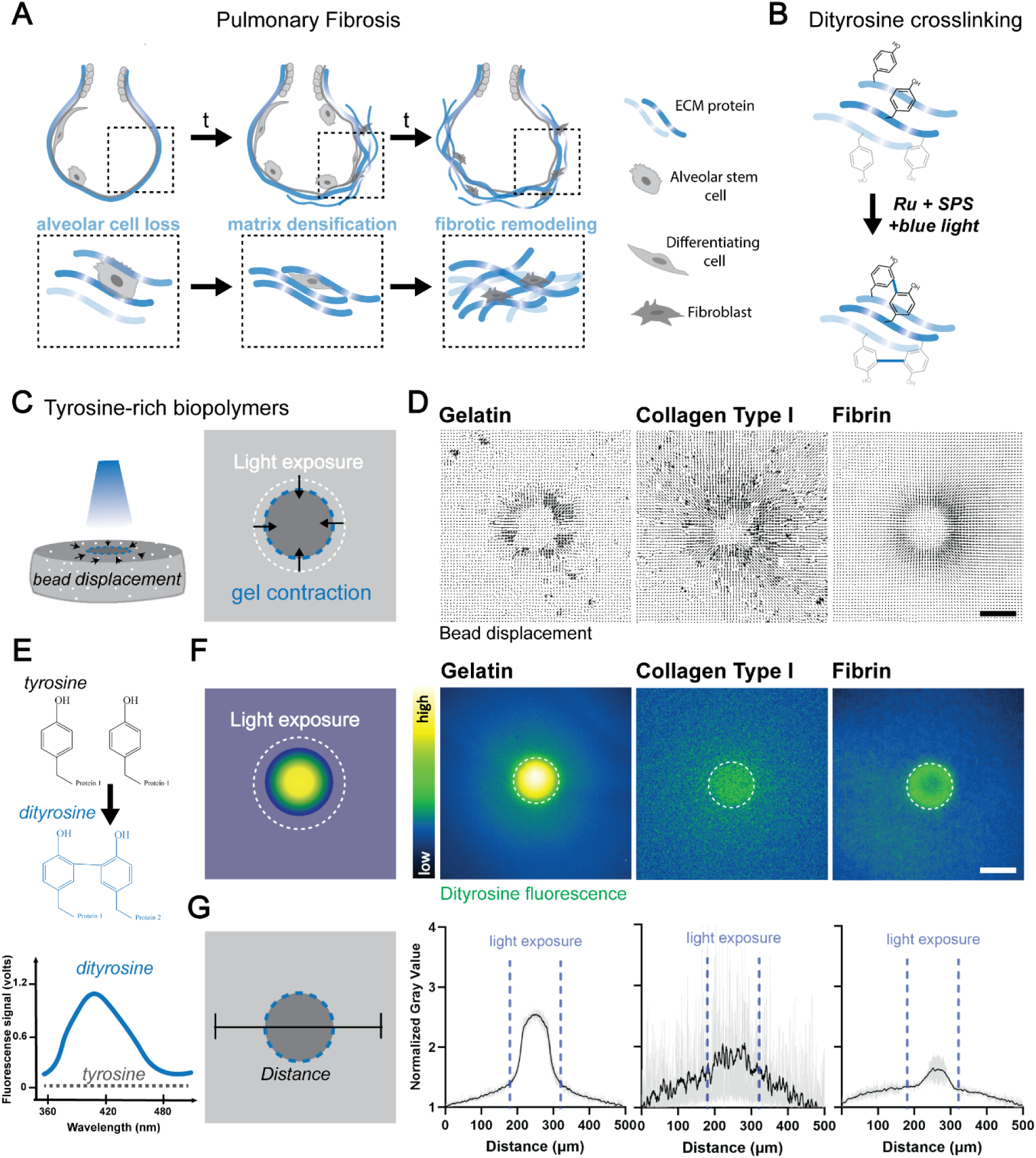
Dityrosine photochemistry enables local crosslinking of ECM proteins *in vitro*. **A**. Schematic illustrating the evolution of pulmonary fibrosis within alveoli initiated by alveolar injury and alveolar cell loss, leading to fibrotic ECM protein deposition, densification, and crosslinking that result in fibrotic remodeling. **B.** Schematic illustrating blue light-mediated crosslinking of dityrosine-rich ECM proteins to mimic ECM densification and crosslinking. **C.** Schematic illustrating local crosslinking of tyrosine-rich biopolymers using focused light and particle image velocimetry (PIV) to visualize the displacement of embedded fluorescent beads upon crosslinking. **D.** Representative PIV plots of bead displacement of gelatin (150 mg/mL), collagen type I (6 mg/mL) and fibrin (5 mg/mL) hydrogels upon blue light exposure (scale bar 100 µm). **E.** Schematic illustrating the formation of dityrosine bonds upon blue light exposure and associated relative fluorescence signals with increased dityrosine fluorescence at 420 nm. **F.** Schematic and representative heat maps of dityrosine fluorescence upon local crosslinking of gelatin, collagen type I and fibrin hydrogels (scale bar 100 µm). **G.** Quantification of normalized pixel intensity of dityrosine fluorescence upon local crosslinking of gelatin, collagen type I, and fibrin hydrogels (mean ± s.e.m for 3 hydrogels per group).

A useful property of dityrosine is that it emits ∼420 nm light under 325 nm excitation ^47,48^ due to the proximity of the two aromatic tyrosine rings, thereby enabling semi-quantitative assessment of the cross-linking reaction (Figure 1E). Across all three hydrogels, dityrosine fluorescence locally increased in response to light and confirmed spatial control over the crosslinking reaction (Figure 1F). Exposure to other wavelengths (e.g., far red light or blue light only (no photo-initiator) did not induce dityrosine fluorescence (Supplementary Fig. 3). Quantification of normalized fluorescence intensities across the ROIs showed an overall fold increase ranging from 1.5 (gelatin), to 1.0 (collagen), and to 0.5 (fibrin) (Figure 1G). Assessment of hydrogel mechanical properties before and after dityrosine crosslinking showed a 1.5-fold increase in storage modulus for gelatin, 9-fold increase for collagen, and 4-fold increase for fibrin gels (Supplementary Fig. 4), likely due to the differences in polymer concentration and tyrosine content. To illustrate the influence of tyrosine content on the formation of dityrosine crosslink density and resulting hydrogel mechanics, we used identical light and photo-sensitizer conditions and compared porcine gelatin (2.6% tyrosine, 150 mg/mL) with bovine gelatin (1% tyrosine, 150 mg/mL). We observed a significant increase in dityrosine fluorescence and storage modulus within the tyrosine-rich porcine gelatin hydrogels (Supplementary Fig. 5), confirming that there is a direct link between tyrosine residue concentration and formation of dityrosine crosslinks ^49^. Although other factors such as polymer fiber length and light scattering may affect the dityrosine fluorescence, these measurements support the use of dityrosine fluorescence as a reliable probe and reproducible technique to assess dityrosine crosslinking density ^47,50^. We further found that varying the concentration and pre-incubation time with the photo-initiator (Ru) provides additional means to modulate the dityrosine crosslinking density and mechanical properties (Supplementary Fig 6). Nano-indentation of gelatin hydrogels confirmed a 1.5-fold increase in Young’s modulus when photo-crosslinked (Supplementary Fig 7). These findings highlight that local blue light exposure enables dityrosine crosslinking of ECM-derived polymer hydrogels to spatiotemporally modulate protein crosslinking and substrate mechanics.

### Local dityrosine crosslinking of ECM proteins within *in situ* lung tissue

Previous work seeking to increase matrix mechanics without changing protein composition demonstrated the potential for dityrosine crosslinking in hydrogels derived from decellularized lung ECM ^51^. Thus, we next sought to apply dityrosine photo-crosslinking *in situ* to precision cut lung slices (PCLS) that maintain the native architecture and composition of alveolar ECM (Figure 2A) ^52,53^. Using 300 µm thick murine lung tissue slices, we observed an increase in dityrosine fluorescence in response to light (Figure 2B). Overall, there was a higher level of background fluorescence when compared to purified biopolymer hydrogels (Figure 1F), presumably due to the pre-existence of dityrosine bonds and general autofluorescence of tissue. Semi-quantitative analysis of dityrosine fluorescence showed a 1.5-fold increase in the region that was exposed to light, indicating successful and spatially defined formation of dityrosine bonds. Given that PCLS from human donor-derived tissue has become a valuable tool towards studying disease mechanisms and drug testing ^54^, we also photo-crosslinked human PCLS (Figure 2C). Human tissue was derived from a healthy donor and dityrosine crosslinking performed within 24 hours after harvesting. Dityrosine fluorescence was also increased 1.5-fold in the photo-crosslinked region when compared to adjacent non-exposed tissue despite an overall higher background fluorescence. Noting that ECM densification and fibrotic remodeling occur in many tissues, we confirmed a similar level of spatial control over dityrosine crosslink formation in skin and liver (Supplementary Fig. 8).

**Figure 2:**
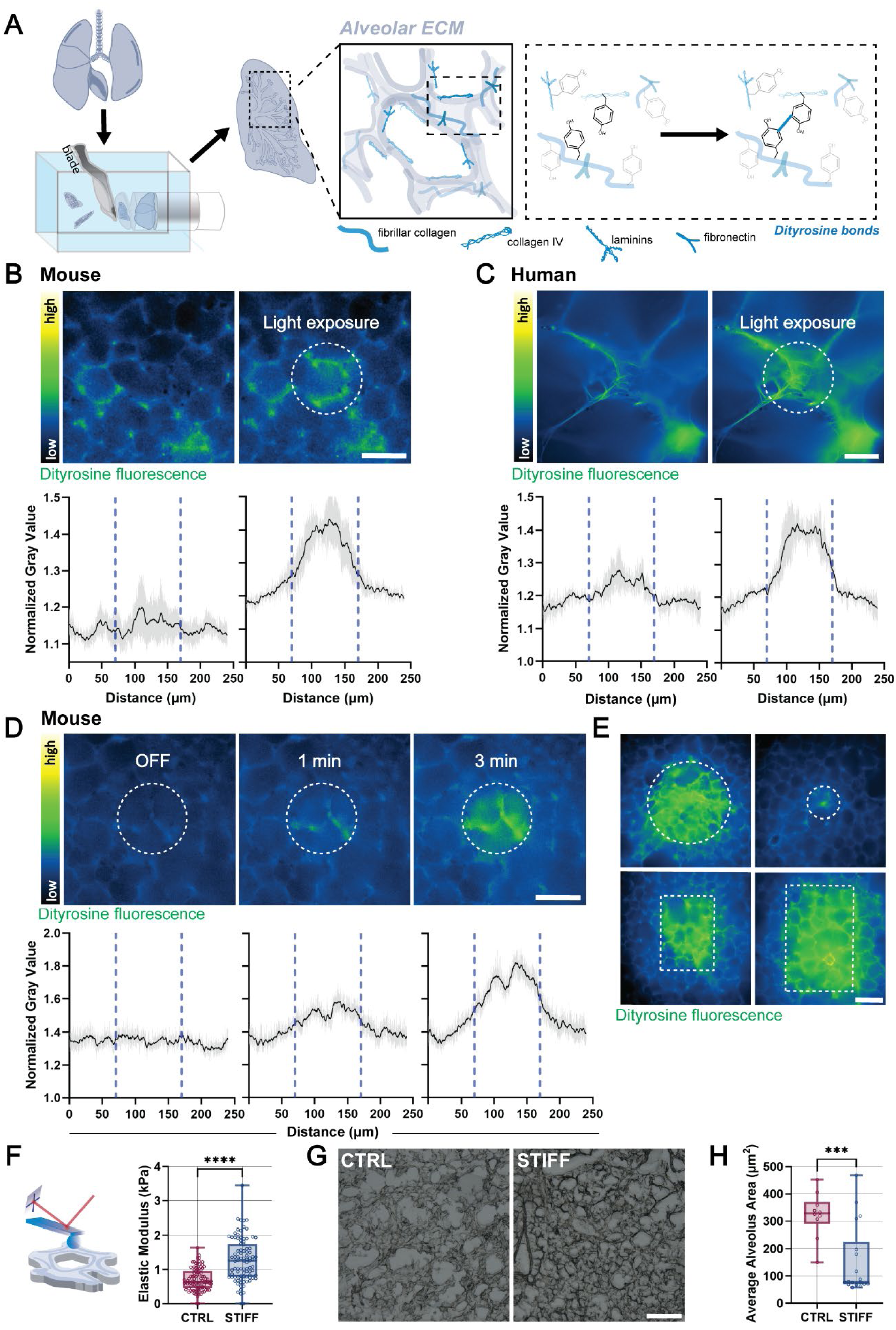
Dityrosine photochemistry enables local crosslinking of ECM proteins in *ex vivo* tissues. **A.** Schematic illustrating the preparation of precision cut lung slices (PCLS) and subsequent blue-light mediated crosslinking of lung alveolar ECM proteins. **B.** Representative heat-maps and quantification of dityrosine fluorescence before and after blue light exposure of murine PCLS (mean ± s.e.m of 3 PCLS, scale bar 50 µm). **C.** Representative heat-maps and quantification of dityrosine fluorescence before and after blue light exposure of human PCLS (mean ± s.e.m of 3 PCLS, scale bar 100 µm). **D.** Representative heat-maps and quantification of dityrosine fluorescence of murine PCLS before (OFF) and after blue light exposure for 1 min and 3 min (mean ± s.e.m of 1 PCLS, scale bar 50 µm). **E.** Representative heat maps of dityrosine fluorescence of photo-crosslinked murine PCLS with various region-of-interests (scale bar 100 µm). **F.** Schematic illustrating atomic force microscopy (AFM) indentation of PCLS parenchyma and quantification of elastic moduli (n = 30 measurements from 3 PCLS from 1 mouse, ****p≤0.0001, two-tailed unpaired Students *t*-test with Welch’s correction. **G.** Representative brightfield images of murine PCLS before (Control, CTRL) and after (stiffened, STIFF) photo-crosslinking (0.13 mM Ru, 20 mW/cm^2^ for 3 minutes, scale bar 250 µm). **H.** Quantification of alveolar area of murine PCLS, mean ± s.d., ***p<0.001, two-tailed unpaired Students *t*-test with Welch’s correction.

Using mouse-derived lung tissue, we then asked how exposure time and photoinitiator concentration contributes to dityrosine crosslinking efficiency. Within one minute of light exposure, small areas of dityrosine fluorescence within the photo-crosslinked region were observed which then extended to fill the entire region by three minutes (Figure 2D). Quantification of dityrosine fluorescence confirmed the 1.25-fold increase within one minute and a 1.40-fold increase when light was continued for three minutes. Modulating the photoinitiator concentration was also assessed as an additional mean to tune dityrosine photo-crosslinking (Supplementary Figure 9). Although increasing the ruthenium concentration from 0.13 mM to 0.26 mM increased dityrosine fluorescence, it also enhances background fluorescence. No dityrosine fluorescence was observed when exposing lung tissue to far red light (Supplementary Figure 10). Given that photo-induced crosslinking enables spatial control over the exposed regions, different sizes of circles and rectangular shapes were also investigated (Figure 2E). Atomic force microscopy indentation was used to measure the local elastic modulus of the lung alveolar ECM before and after dityrosine crosslinking (Figure 2F). We observed an approximate 2-fold increase in tissue modulus from 0.7 ± 0.3 kPa to 1.3 ± 0.6 kPa after in response to light, indicating the formation of functional dityrosine crosslinks and similar mechanical properties that have been measured in *in vivo* murine lung fibrosis models ^55^. Notably, quantification of the average alveolar area in brightfield images of lung tissue (Figure 2G) showed a decrease in alveolar space in stiffened slices (Figure 2H), suggesting densification of the alveolar ECM after dityrosine photo-crosslinking similar to what was observed within ECM-derived polymers (Supplementary Fig. 2). Taken together, these results indicate that blue light enables local stiffening of *in situ* tissue through dityrosine photo-crosslinking of ECM proteins, which is also generalizable to other tissue types.

### Local ECM stiffening directs alveolar epithelial cell function

Having established that dityrosine crosslinking induces lung ECM stiffening similar to early fibrotic remodeling, we next sought to understand whether this change in ECM mechanics alters alveolar epithelial cell differentiation. Cell viability remained high (> 85%) for at least 5 days of PCLS *ex vivo* culture using standard media (DMEM/F12 + 10% fetal bovine serum, Supplementary Figure 11), which is consistent with previous studies ^56–58^. We then confirmed that dityrosine photo-crosslinking (STIFF) has no impact on cell viability when compared to control regions (CTRL) (Figure 3A). To assess whether ECM stiffening mediates alveolar epithelial cell fate within *ex vivo* tissues, we used lineage-tracing of cuboidal AT2 cells, the progenitor/stem cells within the alveoli. Tamoxifen-inducible Sftpc^CreERT2^; ROSA^mTmG^ transgenic mice have widely been used in the field, in which all AT2 cells and their progeny express GFP (Figure 3B) ^14,16^. Indeed, immunostaining of lysosomal associated membrane protein 3 (LAMP3) and podoplanin (PDPN), established markers of AT2 cells and AT1 cells, respectively, showed high levels of LAMP3 but low PDPN staining in freshly isolated PCLS (Supplementary Figure 12), indicating successful AT2 labeling in SftpcCreERT2;ROSA^mTmG^ mice. In CTRL tissues without ECM stiffening, immunostaining for PDPN of GFP-positive cells showed an almost 10x fold increase over the 5 days culture period (Figure 3C, Supplementary Figure 13) with reduced levels of LAMP3 (Supplementary Figure 14), suggesting a steady increase in AT2-to-AT1 differentiation during *ex vivo* culture^53,59^.

**Figure 3:**
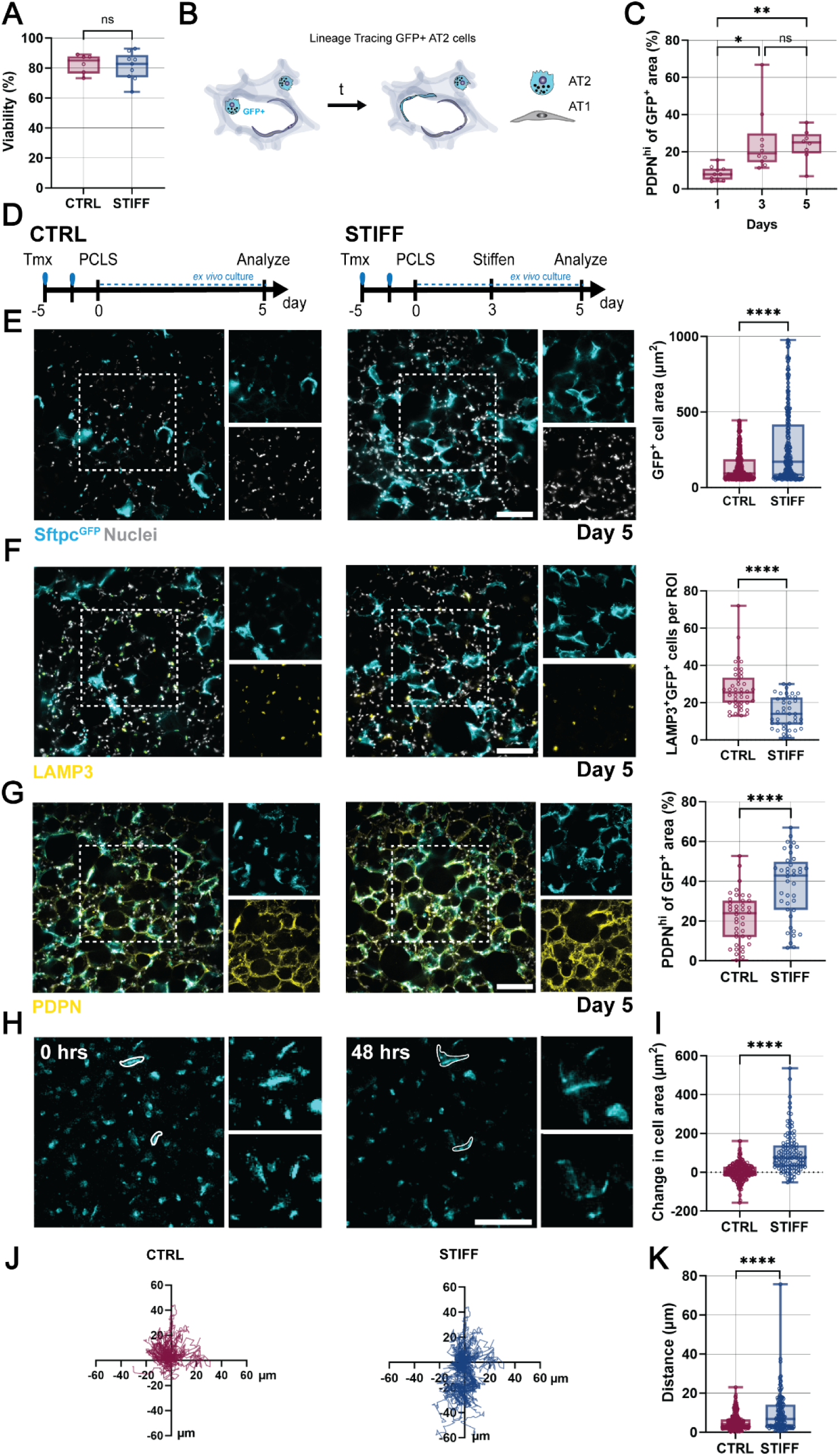
ECM stiffening of ex vivo tissue directs AT2 cell differentiation. **A.** Quantification of PCLS cell viability (Hoechst and Ethidium homodimer-1) in control (CTRL) and stiffened (STIFF) regions at day 2 after blue light exposure (n= 3 images per group from 1 mouse, ns = not significant by two-tailed unpaired Student’s *t*-test). **B.** Schematic illustrating the mouse model to lineage-trace surfactant protein C (SFTPC) expressing type 2 alveolar epithelial (AT2) cells and their progeny using a tamoxifen-inducible SftpcCreERT2;mTmG mice. **C.** Quantification of AT1 specific podoplanin (PDPN) positive cells (high expression) in murine PCLS up to day 5 of *ex vivo* culture (n = 10 images per timepoint from 1 mouse, mean ± s.d.,\*\**p* ≤ 0.01, \**p* ≤ 0.05, ns = not significant by one-way ANOVA with Welch’s correction for multiple comparisons. **D.** Schematic illustrating the experimental timeline including two tamoxifen injection of SftpcCreERT2;mTmG mice 5 days prior preparation of PCLS without (CTRL) or with blue light-mediated ECM stiffening at day 3 (STIFF) and analysis after day 5 of culture. **E.** Representative fluorescent images and quantification of lineage-traced GFP^+^ cell area in CTRL and STIFF regions at day 5 (n = 392 cells (CTRL), n = 276 cells (STIFF) from 2 mice, mean ± s.d., \*\*\*\**p* ≤ 0.0001 by two-tailed unpaired Students *t*-test with Welch’s correction, scale bar 100 µm). **F.** Representative fluorescent images and quantification of LAMP3^+^GFP^+^ cells in CTRL and STIFF regions at day 5 (n = 40 images per group (CTRL, STIFF) from 5 mice, mean ± s.d., \*\*\*\**p* ≤ 0.0001 by two-tailed unpaired Students *t*-test with Welch’s correction, scale bar 100 µm) **G.** Representative fluorescent images and quantification of PDPN^hi^ of GFP^+^ area staining in CTRL and STIFF regions at day 5 (n = 40 images per group (CTRL, STIFF) from 5 mice, mean ± s.d., \*\*\*\**p* ≤ 0.0001 by two-tailed unpaired Students *t*-test with Welch’s correction, scale bar 100 µm). **H.** Representative live cell fluorescent images of Sftpc lineage traced cells in STIFF regions at 0 hours and 48 hours **I.** Quantification of change in cell area in CTRL and STIFF regions of PCLS over 48 hours (n = 240 cells (CTRL), n = 120 cells (STIFF) from 3 mice, \*\*\*\**p* ≤ 0.0001 by two-tailed unpaired Students *t*-test with Welch’s correction). **J.** Migration plots of Sftpc lineage traced cells in CTRL and STIFF regions over 48 hours (n = 60 cells per group). **K.** Quantification of the distance of single cells migrated within 48 hours in CTRL and STIFF regions (n = 120 cells (CTRL) and 300 cells (STIFF) from 2 regions per group from 3 mice, \*\*\*\**p* ≤ 0.0001 by two-tailed unpaired Student’s *t*-test.

After identifying this baseline of alveolar epithelial cell differentiation, we next explored the effect of ECM stiffening. At day 3 after tissue isolation, each tissue slice was split into two regions - either protected from light (i.e., non-crosslinked (CTRL)) or crosslinked with blue light (STIFF) followed by assessment on day 5 (Figure 3D). Elongated morphologies of initially cuboidal GFP^+^ alveolar epithelial cells were observed in both CTRL and STIFF groups with a significant increase in average cell spread area upon ECM stiffening (Figure 3E), suggesting an increase in AT2 cell differentiation ^60^. Thus, we stained for LAMP3, which was observed in both CTRL and STIFF groups but significantly reduced in response to ECM stiffening (STIFF, Figure 3F). This reduction in AT2 cells in STIFF lung tissues was confirmed with an additional AT2 marker (pro-surfactant protein C, proSPC, Supplementary Figure 15). In contrast, expression of PDPN in GFP^+^ alveolar epithelial cells was significantly increased upon ECM stiffening (Figure 3G), suggesting an increase in AT2 differentiation. Given that alveolar epithelial cell motility is functionally important in alveologenesis and response to lung injury ^61,62^, live cell imaging and analysis of migratory behavior in response to ECM stiffening were also investigated (Figure 3H). Trends in enhanced GFP^+^ cell spreading in response to ECM stiffening were confirmed under live cell imaging compared to standard culture (Figure 3I). Tracking analysis of GFP^+^ cells showed an increase in cell migration distance over 48 hours following ECM stiffening (STIFF) when compared to non-crosslinked tissue (CTRL, Figure 3J-K, Supplementary Fig. 16, Supplementary Video 1). These results highlight that local ECM stiffening within *ex vivo* lung tissue directs AT2 cell differentiation and migration by providing mechanical signals mimicking the early fibrotic niche.

### ECM stiffening induces AT2 transitional cell states via mechanosensing

Given that ECM stiffening induced GFP^+^ epithelial cell spreading, we next sought to understand whether these changes in ECM mechanics alter cellular mechanosensing. An important signaling pathway during actin cytoskeletal assembly and contractility includes Rho-associated protein kinase (ROCK) ^63^ (Figure 4A). To investigate how actin cytoskeletal assembly regulates AT2 differentiation in response to ECM stiffening, we inhibited ROCK activity with Y27632 (Y27), added directly to PCLS media after ECM stiffening, and refreshed after 24 hours. After 2 days, both cell spread area and PDPN expression were decreased upon ROCK inhibition (Figure 4B-C). Y27 treatment induced no change in LAMP3 expressing GFP^+^ cells (Figure 4C), suggesting that it is not sufficient to rescue AT2 phenotypes. In contrast, activating RhoA/ROCK with the Rho Activator II (CN03) in non-stiffened CTRL samples led to increased PDPN expression (Supplementary Figure 17). Collectively, these results suggest that cell contractility is sufficient for AT2 differentiation in response to ECM stiffening.

**Figure 4:**
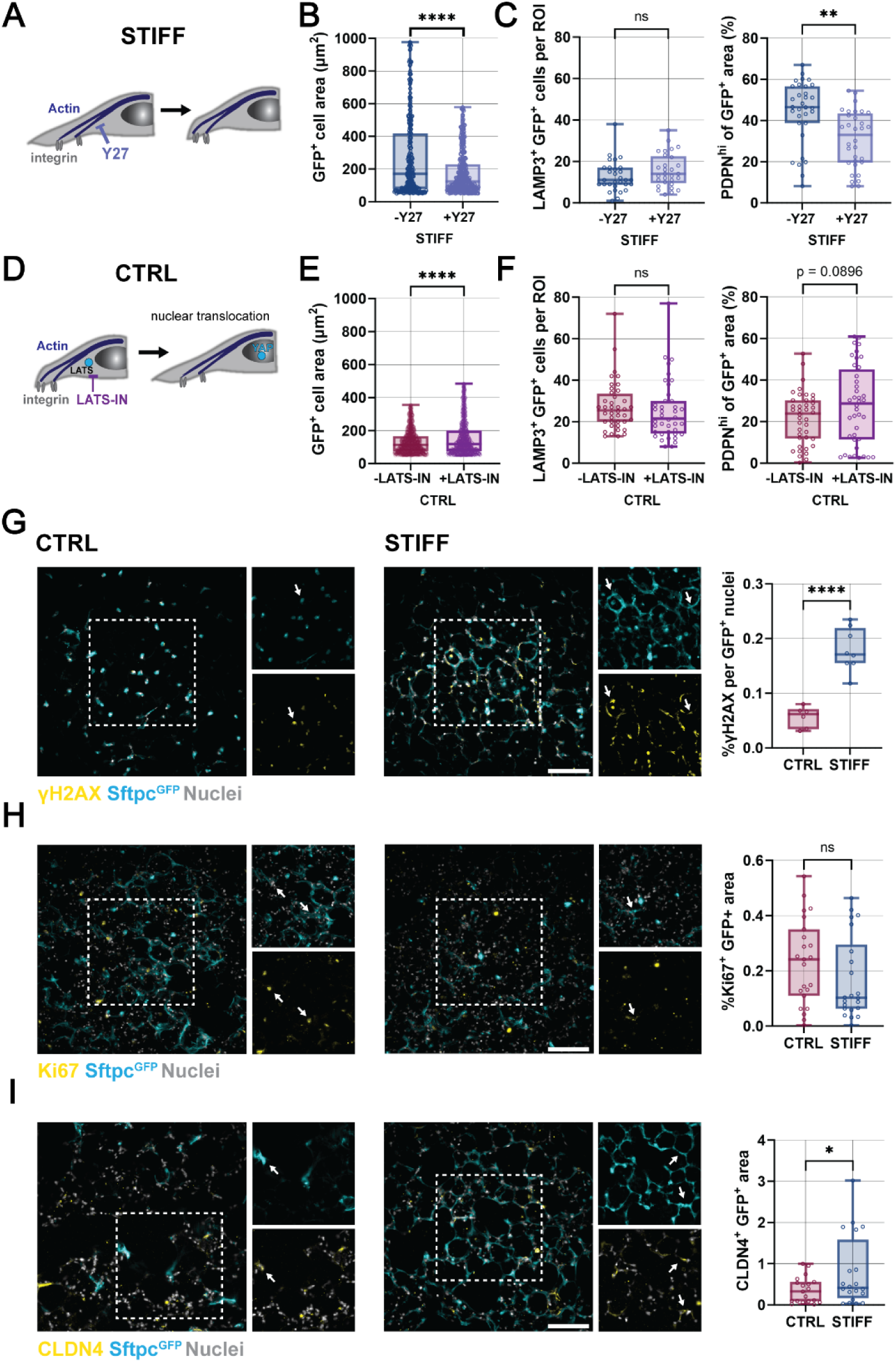
Modulation of mechanosensitive pathways is not sufficient to fully replicate the effects of tissue stiffening. **A.** Schematic illustrating the mechanism of inhibiting actin cytoskeletal tension via addition of Y-27632 (Y27). **B.** Quantification of GFP^+^ cell area in STIFF regions of murine PCLS without and with 10 µM Y-27632 for 2 days (n = 276 cells (STIFF-Y27), n = 400 cells (STIFF +Y27) from 2 mice, mean ± s.d., ****p ≤ 0.0001 and ns = not significant by two-tailed unpaired Student *t*-test with Welch’s correction). **C.** Quantification of LAMP3^+^GFP^+^, and %PDPN^hi^ of GFP^+^ cell area in STIFF regions of murine PCLS without or with 10 µM Y-27632 for 2 days (n = 32 images per group from 4 mice (STIFF-Y27, STIFF +Y27), mean ± s.d., **p ≤ 0.01 and ns = not significant by two-tailed unpaired *t*-test with Welch’s correction). **D.** Schematic illustrating the mechanism of LATS inhibition via addition of LATS-IN-1 to force YAP nuclear translocation. **E.** Quantification of GFP^+^ cell area in CTRL regions of murine PCLS treated without and with 20 µM LATS-IN-1 for 2 days (n = 870 cells (CTRL-LATS-IN), n = 779 cells (CTRL +LATS-IN) per group from two mice, mean ± s.d.,****p ≤ 0.0001 by two-tailed unpaired Student *t*-test with Welch’s correction). **F.** Quantification of LAMP3^+^GFP^+^ cells per ROI and %PDPN^hi^ of GFP^+^ cell area in CTRL regions of murine PCLS without or with 20 µM LATS-IN-1 for 2 days (n = 32 images per group from 4 mice (CTRL-LATS-IN, CTRL +LATS-IN), mean ± s.d., ns = not significant and p as indicated by two-tailed Student’s unpaired *t-*test with Welch’s correction). **G.** Representative images superimposed with GFP^+^ cell mask and quantification of %γH2AX of GFP^+^ nuclei in CTRL and STIFF regions of murine PCLS at day 5 (n = 6 images (CTRL) and 8 images (STIFF) from 2 mice, mean ± s.d., \**p*≤0.05 by two-tailed unpaired Student’s *t*-test with Welch’s correction, scale bar 100 µm) **H.** Representative images and quantification of %Ki67^+^GFP^+^ cells in CTRL and STIFF regions of murine PCLS at day 5 (n = 23 images (CTRL) and 22 images (STIFF) from 5 mice, mean ± s.d., ns = not significant by two-tailed unpaired Student’s *t*-test with Welch’s correction, scale bar 100 µm). **I.** Representative fluorescent images of claudin-4 (CLDN4) immunostaining and quantification of %CLDN4 of GFP^+^ cell area in CTRL and STIFF regions of murine PCLS at day 5 (n = 19 images (CTRL) and 20 images (STIFF) from 5 mice, mean ± s.d., \**p*≤0.05 by two-tailed unpaired Student’s *t*-test with Welch’s correction, scale bar 100 µm).

Similarly, mechanosignaling downstream of the RhoA/ROCK pathway may induce AT2 differentiation. When ROCK is activated, YAP is phosphorylated and sequestered in the nucleus through LATS1/2 inhibition. To assess this, we treated CTRL tissue with large tumor suppressor kinase inhibitor 1 (LATS-IN-1) to biochemically induce nuclear translocation of Yes-associated protein/transcriptional co-activator (YAP/TAZ), which is critical in the transduction of mechanical signals and AT2 cell differentiation ^10,64^ (Figure 4D). Although inducing YAP translocation and activation without ECM stiffening resulted in increased cell spreading (Figure 4E), it did not alter LAMP3 and PDPN expression (Figure 4F). This suggests that YAP activation alone is not sufficient to induce AT1 differentiation.

Recent studies have highlighted that incomplete AT2-to-AT1 differentiation and subsequent accumulation of AT2 transitional cells are associated with fibrotic lesions in both mouse and human lung ^14–16^. The persistence of AT2 transitional cell states may contribute to fibrotic remodeling through mechanisms that include DNA damage and senescence^15,16,65^. To assess whether ECM stiffening prevents complete AT2-to-AT1 differentiation, markers of AT2 transitional cells were assessed. Histone γ-H2AX is one of the most sensitive senescence markers of double-stranded DNA breaks and telomere shortening, and a characteristic marker for AT2 transitional cells ^66^. Two days after ECM stiffening, γ-H2AX showed higher expression in nuclei of GFP^+^ cells in STIFF samples when compared to CTRL samples that were not stiffened (Figure 4G). Even though ECM stiffening increased AT2 senescence, it showed minimal change in Ki67 expression in GFP^+^ cells (Figure 4H) ^15,65^. Claudin-4 (CLDN4) was also used as a characteristic marker for AT2 transitional cells ^15,67^. Two days after ECM stiffening, ∼0.8% of GFP^+^ cells expressed CLDN4 whereas ∼0.4% of GFP^+^ in CTRL samples were CLDN4 positive (Figure 4I). As such, ECM stiffening seems to enhance AT2 transitional cell states with reduced proliferative capacity, similar to what has been observed in fibrotic lesions in mouse and human fibrotic tissue.

### Laminin adhesion regulates AT2 differentiation

Previous studies have shown that fibrotic lesions within the alveoli are characterized by changes in ECM composition and architecture which may impact alveolar epithelial cell function ^2,37^. We first stained for laminin, a major component of the basement membrane and regulator of epithelial cell integrity and proliferation ^68^. Within 2 days after ECM stiffening, pan-laminin staining showed similar architectures in CTRL and STIFF tissues, but a two-fold increase in the projected area of deposited laminin in stiffened regions (Figure 5A). These findings are consistent with previous reports of increased laminin deposition in fibrotic foci ^69^. Several isoforms of the laminin alpha and beta chains (e.g., α1, α3, α5, β4, and α6) have been reported in lung tissue ^4,70^, with laminin α1 (laminin 111 or laminin 1) as a critical regulator of lung development and basement membrane formation ^71^. As cells directly adhere to laminin through integrins such as integrin α6β4 (specific to laminin 111) we next stained for integrin β4, which was similarly expressed in in GFP^+^ cells of CTRL and STIFF samples (Figure 5B). These findings suggest that alveolar epithelial cells maintain adhesion to laminin. To investigate whether this integrin β4 binding to laminin influences AT2 differentiation, we next added a function-blocking integrin β4 antibody (10 µg/mL) directly after ECM stiffening. Blocking integrin β4 did not affect GFP^+^ cell spreading, and higher concentrations of the antibody had no additional effects (Supplementary Figure 18). However, when treated for two days, pan-laminin deposition was reduced for both CTRL and STIFF samples with most significant decrease in STIFF tissue (Figure 5C). This shows that cell-laminin interactions are required for laminin deposition, further suggesting a critical role of alveolar epithelial cells in laminin homeostasis ^72,73^. Interestingly, blocking laminin binding further decreased LAMP3 expression in GFP^+^ cells in both CTRL and STIFF with lowest expression in STIFF + integrin β4 inhibitor treated samples (Figure 5D). In contrast, treatment with integrin β4 antibody had no effect on PDPN expression in CTRL and STIFF tissues, suggesting that AT2 differentiation is independent of integrin β4-mediated adhesion to laminin (Figure 5E).

**Figure 5:**
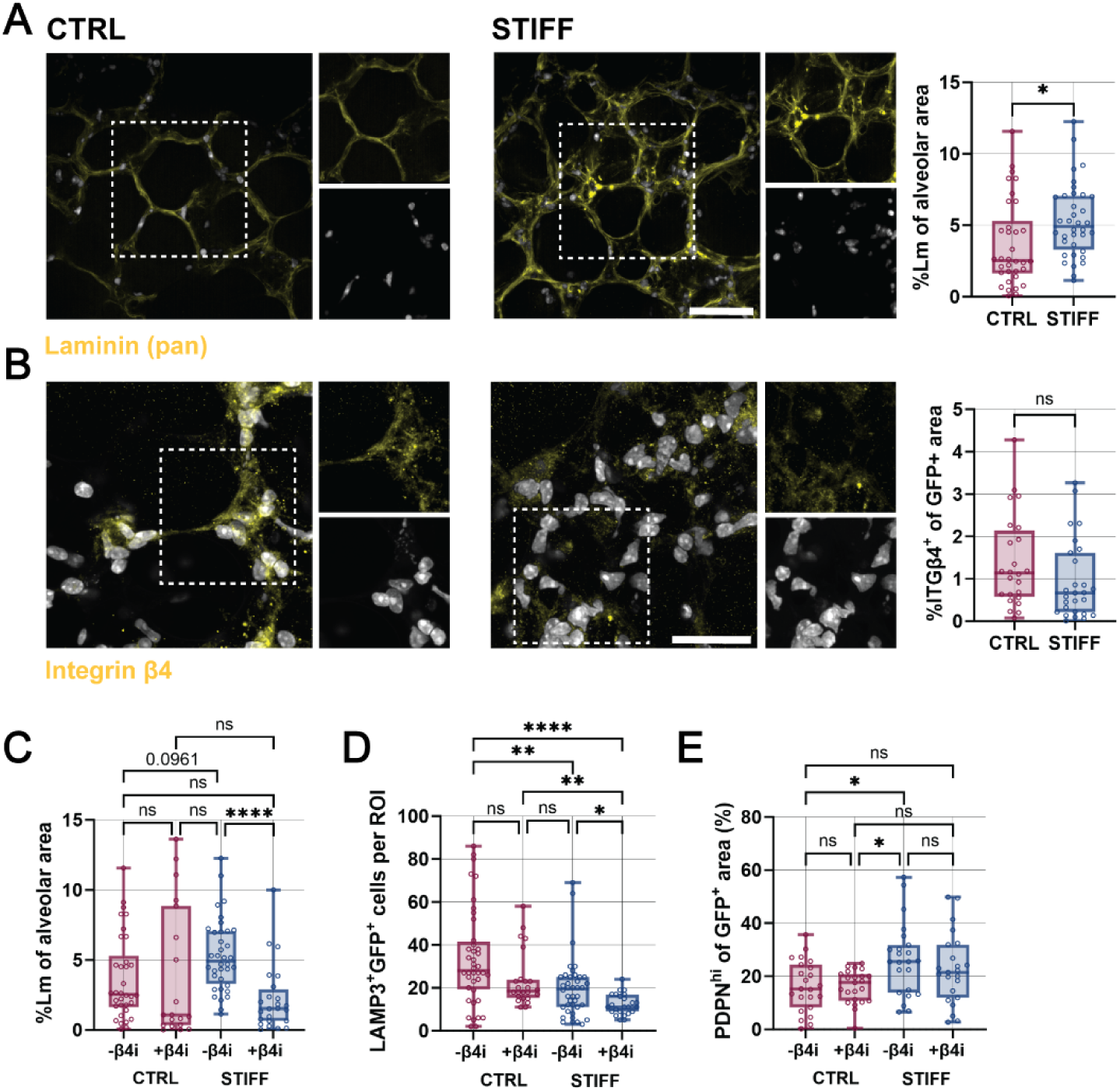
ECM stiffening directs integrin β4-mediated alveolar cell differentiation. **A.** Representative fluorescent images and quantification of pan-laminin (Lm) immunostaining per alveolar area in CTRL and STIFF regions of murine PCLS at day 5 (n = 33 images (CTRL) and 36 images (STIFF) from 5 mice, mean ± s.d., \**p*≤0.05 by two-tailed unpaired Student’s *t*-test with Welch’s correction, scale bar 50 µm). **B.** Representative fluorescent images of integrin β4 (ITGβ4) immunostaining and integrin β4/GFP^+^ area in CTRL and STIFF regions of murine PCLS at day 5 (n = 24 images (CTRL) and 28 images (STIFF) from 5 mice, mean ± s.d., ns = not significant by two-tailed unpaired Student’s *t*-test with Welch’s correction, scale bar 30 µm). **C** Quantification of pan-laminin immunostaining per alveolar area in CTRL and STIFF regions of murine PCLS without or with 10 µg/mL integrin β4 function-perturbing antibody (+β4i) for 2 days (n = 33 images (CTRL) and 36 images (STIFF) from 5 mice, and n = 19 images (CTRL + β4i) and 24 images (STIFF + β4i) from 4 mice, mean ± s.d., ****p ≤ 0.0001, p as indicated, ns = not significant by one-way ANOVA with Tukey’s multiple comparisons test). **D** Quantification of LAMP3^+^GFP^+^ cells per ROI in CTRL and STIFF regions of murine PCLS without or with 10 µg/mL integrin β4 function perturbing antibody (+β4i) for 2 days (days (n = 40 images per group (CTRL, STIFF) from 5 mice, and n = 24 images per group (CTRL + β4i, STIFF + β4i) from 3 mice, , mean ± s.d., ****p ≤ 0.0001, **p ≤ 0.01, *p ≤ 0.05, and ns = not significant by one-way ANOVA with Tukey’s multiple comparisons test. **E** Quantification of PDPN^hi^ of GFP^+^ area in CTRL and STIFF regions of murine PCLS without or with 10 µg/mL integrin β4 function perturbing antibody (+β4i) for 2 days (n = 23 images per group (CTRL, STIFF) from 3 mice, and n = 23 images per group (CTRL + β4i, STIFF + β4i) from 3 mice, mean ± s.d., *p ≤ 0.05 and ns = not significant by one-way ANOVA with Tukey’s multiple comparisons test).

Collectively, these findings suggest that, in response to ECM stiffening, GFP^+^ cells downregulate their interactions with laminin which affects AT2 cell maintenance. This is consistent with reduced AT2 proliferation after ECM stiffening (Figure 4H) and previous studies reporting integrin β4 as a key regulator of human lung epithelial cell proliferation and differentiation^74^.

### AT2 differentiation requires Integrin β1 expression

Previous studies in mouse and human fibrotic lesions demonstrated an increase in aberrant ECM accumulation, such as fibronectin, an ECM protein that is typically secreted by fibroblasts/myofibroblasts in response to injury ^75^. Thus, we next tested whether ECM stiffening influences the deposition and interaction of alveolar epithelial cells with fibronectin. Fibronectin staining was expressed throughout the tissue in both CTRL and STIFF tissue (Figure 6A). However, ECM stiffening significantly increased the deposition of fibronectin compared to CTRL tissue (Figure 6B), with a trend towards higher expression of alpha smooth muscle actin (αSMA), suggesting that myofibroblast may start to get activated (Supplementary Figure 19). A major cellular binding site of fibronectin is its RGD domain which is also involved in cell-mediated ECM remodeling ^76^. Selectively blocking fibronectin binding sites with the monoclonal antibody HFN7.1 (10 µg/mL) reduced GFP^+^ cell spread area, and higher concentrations (5-15 µg/mL) had little additional effect (Supplementary Figure 20). However, no significant difference was observed in both LAMP3 and PDPN expression of GFP^+^ cells, Figure 6C), suggesting that GFP^+^ cell adhesion to the RGD domain of fibronectin has no critical function in AT2 maintenance or differentiation.

**Figure 6:**
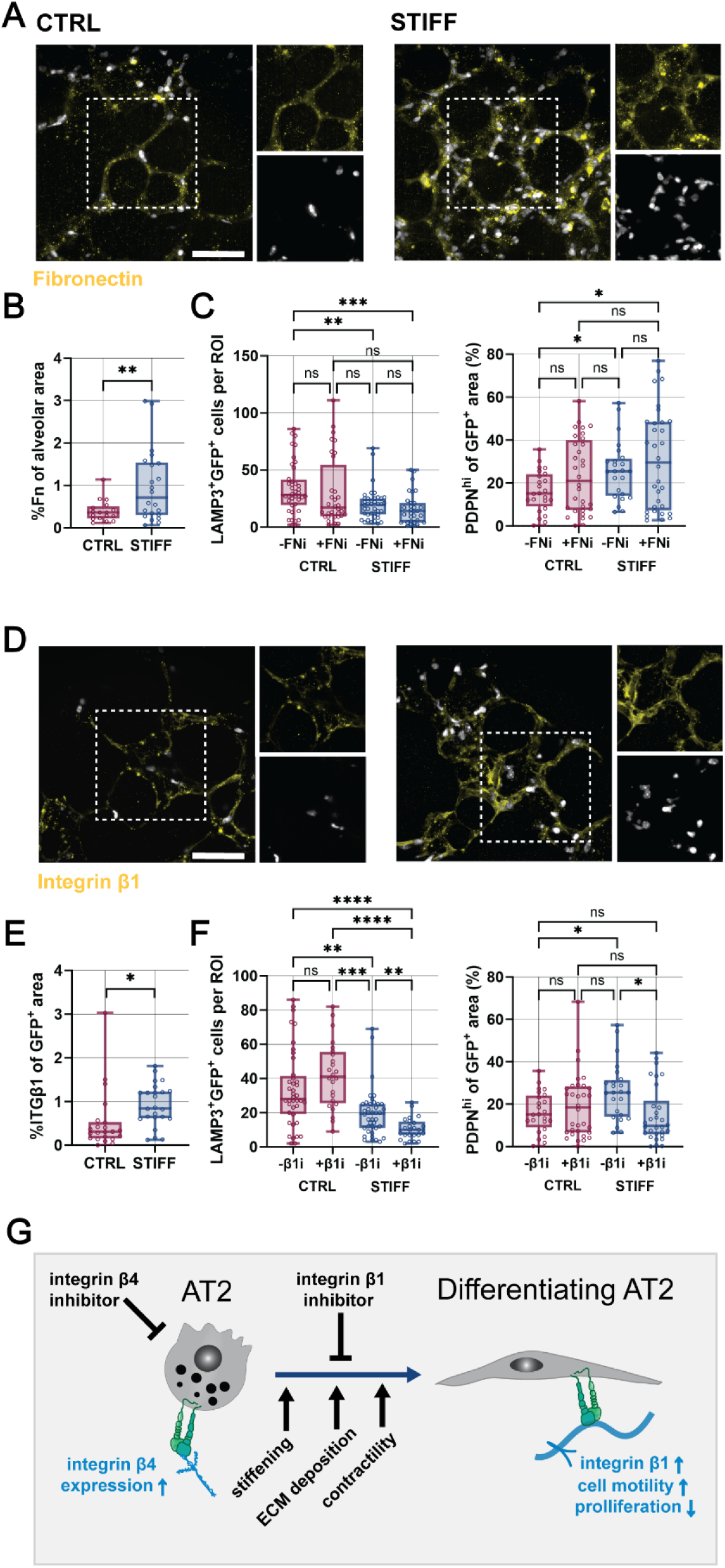
Integrin β1 is required for alveolar epithelial cell differentiation in response to ECM stiffening. **A.** Representative fluorescent images of fibronectin immunostaining in CTRL and STIFF regions of murine PCLS at day 5 (scale bar 50 µm). **B.** Quantification of fibronectin (Fn) expression per alveolar area in CTRL and STIFF regions of murine PCLS (n = 21 images (CTRL) and 22 images (STIFF) from 5 mice at day 5, mean ± s.d., \*\**p* ≤ 0.01 by two-tailed unpaired Student’s *t*-test with Welch’s correction). **C** Quantification of LAMP3^+^GFP^+^ cells in CTRL and STIFF regions of murine PCLS without or with 10 µg/mL fibronectin function perturbing antibody (HFN7.1) for 2 days (n = 40 images per group from 5 mice (CTRL, STIFF), 32 images per group from 4 mice (CTRL+FNi, STIFF+FNi), mean ± s.d., **p < 0.01, ***p < 0.001, ns = not significant by one-way ANOVA with Tukey’s multiple comparisons test and quantification of PDPN^hi^ of GFP^+^ area in CTRL and STIFF regions of murine PCLS without or with 10 µg/mL fibronectin function perturbing antibody (HFN7.1) for 2 days (n = 24 images per group from 3 mice (CTRL, STIFF), 24 images per group from 3 mice (CTRL+FNi, STIFF+FNi), mean ± s.d., *p < 0.05, ns = not significant by one-way ANOVA with Tukey’s multiple comparisons test). **D.** Representative fluorescent images of integrin β1 immunostaining in CTRL and STIFF regions of murine PCLS at day 5 (scale bar 50 µm). **E.** Quantification of integrin β1 (ITGβ1) expression of GFP^+^ cells in CTRL and STIFF regions of murine PCLS (n = 22 images (CTRL) and 24 images (STIFF) from 4 mice at day 5, mean ± s.d., \**p* ≤ 0.05 by two-tailed unpaired Student’s *t*-test with Welch’s correction). **F.** Quantification of LAMP3^+^GFP^+^ cells in CTRL and STIFF regions of murine PCLS without or with 10 µg/mL integrin β1 function perturbing antibody (β1i) for 2 days (n = 40 images per group from 5 mice (CTRL, STIFF), 24 images per group from 3 mice (CTRL+β1i, STIFF+ β1i), mean ± s.d., \*\**p* ≤ 0.01, \*\*\**p* ≤ 0.001, \*\*\*\**p* ≤ 0.0001, ns = not significant by one-way ANOVA with Tukey’s multiple comparisons test). and quantification of PDPN^hi^ of GFP^+^ area in CTRL and STIFF regions of murine PCLS without or with 10 µg/mL integrin β1 function perturbing antibody (β1i) for 2 days (n = 24 images per group from 3 mice (CTRL, STIFF), 32 images per group from 4 mice (CTRL+ β1i, STIFF+ β1i), mean ± s.d., *p < 0.05, ns = not significant by one-way ANOVA with Tukey’s multiple comparisons test). **G.** Schematic illustrating the proposed mechanism of integrin engagement and ECM deposition as a regulator of alveolar epithelial cell differentiation in response to blue light mediated ECM stiffening as a model of early fibrotic remodeling within the alveoli.

Given that cells may also interact with fibronectin through other domains than RGD (i.e. α4β1) ^77–79^ we next stained for integrin β1, which is expressed by AT2 cells ^80^ and is a major integrin for cellular interactions with fibronectin (Figure 6D). Within 2 days after ECM stiffening, integrin β1 was significantly upregulated in GFP^+^ cells when compared to CTRL tissues (Figure 6E). When the binding domain of integrin β1 was blocked with an AIIB2 antibody (10 µg/mL), cell spreading was reduced with minimal additional effect of varying antibody concentration (Supplementary Figure 21), similar to integrin β4 and HFN7.1 perturbation (Supplementary Figure 18 and 20). Notably, blocking integrin β1 in CTRL tissue may be able to rescue AT2 phenotypes, although this trend was not significant (Figure 6F). Still, this is consistent with previous reports showing enhanced AT2 persistence is present in β1 deficient mice ^81^. However, blocking integrin β1 was not sufficient to maintain LAMP3 expression in ECM stiffened samples, suggesting that increased ECM mechanics may override the initial perturbation of integrin β1 engagement. PDPN staining showed minimal changes in CTRL tissue. Notably, blocking integrin β1 induced a significant reduction in PDPN expression in response to ECM stiffening, highlighting that β1 binding is required for AT2 differentiation. Taken together, our findings suggest that alveolar epithelial cell function is influenced by ECM stiffening and that integrin β1 is a critical mediator of AT2 maintenance and differentiation.

### Outlook

Several studies have demonstrated that an increase in ECM mechanics is characteristic of fibrotic remodeling in lung tissue. Often, a focus has been placed on the influence of ECM mechanics on fibroblast spreading and differentiation into aberrant ECM-producing myofibroblasts that drive disease progression. Our findings indicate that local ECM stiffening not only influences ECM deposition, but also the adhesion and differentiation of alveolar epithelial cells. This has not been studied yet, due to the lack of approaches to modulate ECM stiffness *in vivo* or *ex vivo*. Using a dityrosine photochemistry, we developed an *ex vivo* early-stage fibrosis model, which enables on demand local crosslinking and stiffening of the ECM within the native tissue microenvironment. Specifically, we found that ECM stiffening within *ex vivo* lung tissue induces AT2 differentiation that was inhibited when intracellular contractility or integrin β1 engagement was blocked (Figure 6G). Increased γH2AX and CLDN4 expression in these cells further suggests that ECM stiffening increases accumulation of AT2 transitional cells that have been associated with fibrotic lesions in mouse and human lung fibrosis ^15,66^. Thus, our study now demonstrates a direct correlation between increased mechanical forces and impaired AT2 differentiation. Previous work has used *in vitro* stretching devices to study the role of mechanical forces in isolated AT2 differentiation ^82^. However, the *ex vivo* tissue stiffening model developed here will provide means to implement signals of the native microenvironment and other cell types such as endothelial cells ^83,84^ or macrophages ^85^. Equibiaxial stretching devices may further be used to study the interplay of ECM stiffening and cyclic breathing mechanics ^86,87^. Embedding ECM stiffened lung tissue slices into synthetic hydrogels may also be relevant for studies that require long term culture conditions ^88^. Taken together, our work shows that dityrosine crosslinking induces ECM stiffening that recapitulates the initial stages of fibrotic lesions and provides an engineered platform to study fibrosis *ex vivo*.

## Methods

### 2D hydrogel fabrication and photo-crosslinking

Gelatin hydrogels were fabricated by mixing 150 mg porcine or bovine gelatin with 1 mL phosphate buffer saline (PBS), incubated at 40°C until solubilized and then pre-crosslinked between two coverslips for 10-15 minutes at room temperature (RT). Fibrin hydrogels were fabricated by pre-crosslinking 5 mg mL^-1^ fibrinogen (Sigma F8630) hydrogels with 1 Unit mL^-1^ thrombin (Sigma T4648), followed by incubation for 30 minutes at 37°C. Collagen type I hydrogels (Fibricol, Advanced Biomatrix 5133) were fabricated by adjusting a 6 mg mL^-1^ collagen solution to neutral pH with 1 N NaOH, followed by incubation for 30 minutes at 37°C. Pre-crosslinked gelatin, fibrin and collagen hydrogels were then incubated in 20 mM SPS sodium persulfate (SPS) and 0.13-1 mM [RuII(bpy)^3^]^2+^ (Advanced Biomatrix 5248) for 20 minutes, followed by local crosslinking using the blue light laser of a widefield fluorescent microscope (Leica THUNDER DMi8) set at 45% intensity for 1-5 minutes.

### Particle image velocimetry analysis

Gelatin, fibrin, and collagen hydrogels were prepared as described with 0.5 µm-diameter fluorescent beads at 2% vol/vol concentration (Bangs Laboratories, FSPP003). Embedded beads were captured before and after local crosslinking for 3-minute exposure time) of a widefield fluorescent microscope as previously described. The ImageJ plugin particle image velocimetry was used to analyze bead displacement of acquired fluorescent images^89^.

### Dityrosine fluorescence imaging and analysis

Gelatin, fibrin, and collagen hydrogels were locally crosslinked as described previously and dityrosine fluorescence visualized using the UV channel at 25x magnification. Regional changes in pixel intensity were quantified in ImageJ by generating at least nine intensity profile plots throughout the crosslinked regions and averaged for each individual image. Intensity profiles were normalized to minimum value of surrounding area near the crosslinked region of interest to represent fold change in fluorescence.

### Mechanical testing

#### Shear photo-rheology

Pre-cursor solutions of gelatin, fibrin and collagen hydrogels were prepared as described and pre-crosslinked between a cone and plate geometry (20 mm diameter, 54 µm gap size) of a HR 30 Discovery Hybrid Rheometer (TA Instruments). A custom poly(dimethylsiloxane) PDMS (Sylgard 184, Ellsworth Adhesives, 9:1 ratio) with an embedded heating coil allow for ruthenium incubation and hydrogel crosslinking at 37°C for fibrin and collagen hydrogels. Measurements of pre-crosslinked and light-induced crosslinked hydrogels were performed by oscillatory time sweeps (0.5 Hz and 0.5% strain).

#### Atomic Force Microscopy (AFM)-guided nanoindentation

Pre-crosslinked and light-crosslinked gelatin hydrogels were prepared as described and AFM-nanomechanical mapping performed using a HYDRA6V-200NG (AppNano) probe and glass microspherical tip (Fisher) (R ≈ 6.79, k ≈ 0.0308 N/m) via a Nanosurf FlexAFM system. Indentations were performed nine times at three distinct regions for a total of 27 points. Force-displacement curves were fit to the Hertz model, assuming a Poisson’s ratio of 0.5 ^90^.

For lung tissue, PCLS samples were embedded in optimal cutting temperature medium and cryo-sectioned into 20-µm-thick slices via the Kawamoto’s film method ^91,92^. AFM-nanoindentation was performed at 10 μm/s z-piezo displacement rate up to ≈ 1 μm indentation depth using a microspherical tip (*R* ≈ 5 µm, *k* ≈ 0.03 N/m, HQ:CSC38/tipless/Cr-Au, cantilever B, NanoAndMore) and a Dimension Icon AFM (BrukerNano) in 1× PBS. To account for spatial heterogeneity, indentation was performed at least 30 randomly selected indentation locations. The effective indentation modulus was calculated by via the finite thickness-corrected Hertz model, assuming Poisson’s ratios of 0.45 ^93^.

### Human Tissue

Normal lungs from deceased individuals were de-identified and obtained from Gift of Life Michigan. Tissue samples were deemed IRB exempt and approved for research use by the University of Michigan Institutional Review Board (IRB).

### Mouse lineage tracing

SftpcCreERT2;mTmG mice were generated by crossing Sftpctm1(cre/ERT2)Blh(Sftpc-CreERT2) with Rosa26-mTmG (mTmG) mice. SftpcCreERT2 mice^94^ were obtained from Dr. Harold Chapman. Rosa26-mTmG (abbreviated mTmG, 007576) were obtained from Jackson Laboratories. All animal husbandry and experiments were approved by the Unit for Laboratory Animal Medicine (ULAM) at the University of Michigan (IACUC #PRO00012071). Lineage tracing was initiated in 6-8 week old SftpcCreERT2;mTmG mice by 100 µL intra-peritoneal injection of tamoxifen in corn oil (Sigma T5648, 20 mg/mL) six and three days prior euthanasia. Both sexes were used for this study.

### Precision-cut lung slice (PCLS) preparation and culture

7-9 week-old mice were euthanized via CO2 inhalation, followed by flushing the lungs with sterile phosphate-buffered saline (PBS) via the heart and cannulation of the lung with a catheter in the anterior wall of the trachea, right above the cricoid cartilage. Next, 1.25 mL of 37°C 2% low-melting agarose in sterile PBS (Sigma 39346-81-1) was injected to inflate both lungs, followed by ligating the trachea with thread to retain the agarose inside the lungs and incubation of the lungs at 4°C for 10 minutes before excision. The excised lungs were then washed with sterile cold PBS, and each separated lung lobe was cut at 300 µm thick using an automated vibratome (VF-210-0Z Compresstome, Precisionary Instruments). Slices were obtained from the middle 2/3rds of the lobe to ensure similar sized slices. PCLS were collected in a 24 well plate (2 slices/well) and cultured in 1.5 mL of DMEM/F12 culture plus 10% fetal bovine serum (FBS) at 37°C and 5% (volume/volume) CO2. Culture medium was changed once 2 hours post-sectioning and then every day of culture for up to 5 days.

### Photo-crosslinking of PCLS

Following 3 days of *ex vivo* culture, PCLS were washed once with PBS, incubated in 20 mM sodium persulfate (SPS) and 0.13-1 mM [RuII(bpy)^3^]^2+^ (Advanced Biomatrix 5248) for 20 minutes at 37°C for 20 minutes in customized poly(dimethylsiloxane) PDMS (Sylgard 184, Ellsworth Adhesives, 9:1 ratio) molds. Local crosslinking was performed using a mylar photomask film (Fine Line Imaging) and blue light (400-500 nm, Omnicure S1500) for 3 min at 25 mW cm^−2^. PCLS were then washed once with PBS and culture continued in DMEM/F12 plus 10% FBS for an additional 2 days prior fixation in 4% paraformaldehyde. The same procedure was used for photo crosslinking of murine skin and liver tissue slices.

### PCLS dityrosine fluorescence and alveolar area imaging and analysis

Dityrosine fluorescence imaging and quantification were performed as described for gelatin hydrogels. Phase contrast images of CTRL and STIFF samples were used to measures the change in airspace area via thresholding of the empty spaces in ImageJ.

### PCLS viability assay

PCLS were first incubated in PBS containing Hoechst (1:1000, Sigma 62249) for 20 minutes at 37°C, followed by an additional 20 minutes at room temperature after adding ethidium homodimer-1 (ETHD-1, 4 µM, Invitrogen E1169). Viability was quantified from 200 µm stacks acquired using a Leica DMi8 THUNDER microscope and reported as the ratio of live cells (Hoechst^+^/ETHD^-^) and the total number of cells.

### PCLS small molecule inhibition

To perturb cell-ECM interactions, PCLS were cultured in DMEM/F12 plus 10% FBS for 3 days, prior adding monoclonal antibodies against fibronectin (10 μg ml^−1^ HFN7.1, Developmental Studies Hybridoma Bank), integrin β1 (10 μg ml^−1^ AIIB2, Developmental Studies Hybridoma Bank), or integrin β4 clone ASC-8 (10 μg ml^−1^, Millipore MAB2059Z), and media refreshed daily. Cell mechanosensing was perturbed using 10 μM ROCK inhibitor (Y-27632, Tocris 1254251), 20 µM LATS-IN (Fisher 50-225-9251), or 5 μg/mL Rho activator II (Cytoskeleton Inc, CN03).

### PCLS immunofluorescence staining

PCLS were fixated with 4% paraformaldehyde for 1 hour at room temperature, washed with PBS, followed by incubation in permeabilization solution (0.2% Triton-X, 10 wt% sucrose) for 45 minutes at 4°C and then blocked in PBS containing 2% BSA 0.1% Triton-X, and 5% horse serum for 2 hours at room temperature. No permeabilization was performed or ECM protein immunostaining to minimize intracellular staining. Primary antibodies were diluted in blocking solution (2% BSA 0.1% Triton-X, and 5% horse serum) and PCLS, incubated overnight at 4 °C, washed three times with PBS, followed by incubation in secondary antibodies and Hoechst staining (1:1000) for 30 minutes at room temperature. Antibodies and dilutions included anti-LAMP3 (1:100; Eurobio DDX0192P), anti-PDPN (1:100, DHSB Q62011), anti-proSPC (Millipore AB3786), anti-HopX (1:200, Santa Cruz sc-398703), anti-ki67 (1:1000, Invitrogen 14-5698-82), anti-claudin 4 (1:100, Invitrogen 36-4800), anti-laminin (1:250, Abcam 11575), anti-integrin β4 (1:100, Abcam ab236251), anti-fibronectin (1:100, Thermofisher 15613- 1-AP), anti-integrin β1 (1:100, Invitrogen PA5-78028), anti-αSMA (1:1000, Abcam ab7817), Alexa Fluor-647 IgG H&L (1:500; Thermofisher A-21451/A-21244/A-48265/A-21235).

### Imaging and quantification

All images were acquired on a Leica DMi8 THUNDER widefield microscope, integrin β4 images were taken on a Zeiss LSM800 confocal microscope with a 63x oil immersion objective. For ECM markers (fibronectin, laminin) and α-SMA immunostaining analysis, a mask of the tdTomato channel of the alveolar area was superimposed with a mask of each respective stain, then thresholded and quantified using ImageJ. For all other remaining immunostaining markers, a mask of the Sftpc^GFP^ cell area was superimposed with a mask of each stain, then thresholded and quantified using ImageJ. For ki67 and γH2AX immunostaining analysis, a mask of both Hoechst and GFP+ area was superimposed with each stain to quantify number of stained GFP+ nuclei.

### Time-lapse imaging of PCLS and quantification

PCLS were prepared and stiffened as described, placed within a 25 mm diameter PDMS mold, and 5 wt% methacrylated hyaluronic acid (meHA) hydrogel pipetted around the tissue and crosslinked with UV light for 3 minutes (3 mW/cm^2^) to reduce movement. After adding 1 mL of culture medium, constructs were placed within a pre-equilibrated and humidified stage-top environmental control chamber (TOKAI Hit), with controlled temperature (37°C), humidity (95%), and gas concentration (5% CO2). 400 µm thick z stacks images were taken every 20 minutes within CTRL and STIFF regions for 48 hour. Images were analyzed using ImageJ by individually tracking at least 30 cells per group with the Manual Tracking plugin. Individual cell trajectories, distance traveled, and velocity were derived from x- and y-coordinates per timepoint obtained from manual tracking.

### Statistical Analysis

Statistical analyses were performed using GraphPad Prism software (version 9&10). Statistical comparisons between two experimental groups within the slices were performed using un-paired two-tailed Student’s t-tests with Welch’s correction and comparisons among more groups were performed using one-way ANOVA. All experiments were repeated as described in the text. For representative immunofluorescence images, at least four biological repeats of all experiments were performed with similar results.

## Acknowledgments

This work was partially supported by funding from the NIH (R00-HL151670 to C.L, NIH T32 GM145304 to D.W.A, NHLBI T32 HL007749 to M.L.T), the American Lung Association (IA-939940 to C.L.), the David and Lucile Packard Foundation (to C.L), and the National Science Foundation (NSF CMMI-1751898 to L.H.). The authors also thank Steven Huang for his assistance with human tissue sample collection.

## References

1. Hackett, T. L. & Osei, E. T. Modeling Extracellular Matrix-Cell Interactions in Lung Repair and Chronic Disease. Cells 10, 2145 (2021).

2. Burgstaller, G. et al. The instructive extracellular matrix of the lung: basic composition and alterations in chronic lung disease. European Respiratory Journal 50, 1601805 (2017).

3. Naba, A., et al. The Matrisome: In Silico Definition and In Vivo Characterization by Proteomics of Normal and Tumor Extracellular Matrices. Molecular & Cellular Proteomics 11, M111.014647 (2012).

4. Balestrini, J. L. & Niklason, L. E. Extracellular matrix as a driver for lung regeneration. Ann Biomed Eng 43, 568 (2015).

5. Waters, C. M., Roan, E. & Navajas, D. Mechanobiology in Lung Epithelial Cells: Measurements, Perturbations, and Responses. Compr Physiol 2, 1 (2012).

6. Zhou, Y. et al. Extracellular matrix in lung development, homeostasis and disease. Matrix Biology 73, 77–104 (2018).

7. Martinez, F. J. et al. Idiopathic pulmonary fibrosis. Nat Rev Dis Primers 3, 17074 (2017).

8. Larsen, B. T. Usual interstitial pneumonia: a clinically significant pattern, but not the final word. Modern Pathology 2022 35:5 35, 589–593 (2022).

9. Burgess, C. L. et al. Generation of human alveolar epithelial type I cells from pluripotent stem cells. Cell Stem Cell 31, 657–675.e8 (2024).

10. Shiraishi, K. et al. Biophysical forces mediated by respiration maintain lung alveolar epithelial cell fate. Cell 186, 1478–1492.e15 (2023).

11. Li, J. et al. The Strength of Mechanical Forces Determines the Differentiation of Alveolar Epithelial Cells. Dev Cell 44, 297–312.e5 (2018).

12. Hogan, B. L. M. et al. Repair and Regeneration of the Respiratory System: Complexity, Plasticity, and Mechanisms of Lung Stem Cell Function. Cell Stem Cell 15, 123–138 (2014).

13. Desai, T. J., Brownfield, D. G. & Krasnow, M. A. Alveolar progenitor and stem cells in lung development, renewal and cancer. Nature 507, 190–194 (2014).

14. Jiang, P. et al. Ineffectual Type 2 to Type 1 Alveolar Epithelial Cell Differentiation in Idiopathic Pulmonary Fibrosis: Persistence of the KRT8hi Transitional State. Am J Respir Crit Care Med 201, 1443–1447 (2020).

15. Kobayashi, Y. et al. Persistence of a regeneration-associated, transitional alveolar epithelial cell state in pulmonary fibrosis. Nature Cell Biology 2020 22:8 22, 934–946 (2020).

16. Strunz, M. et al. Alveolar regeneration through a Krt8+ transitional stem cell state that persists in human lung fibrosis. Nat Commun (2020) doi:10.1038/s41467-020-17358-3.

17. Wolters, P. J., Collard, H. R. & Jones, K. D. Pathogenesis of Idiopathic Pulmonary Fibrosis. Annual Review of Pathology: Mechanisms of Disease 9, 157–179 (2014).

18. Herrera, J., Henke, C. A. & Bitterman, P. B. Extracellular matrix as a driver of progressive fibrosis. J Clin Invest 128, 45–53 (2018).

19. Liu, F. et al. Feedback amplification of fibrosis through matrix stiffening and COX-2 suppression. J Cell Biol 190, 693–706 (2010).

20. Tschumperlin, D. J. Matrix, mesenchyme, and mechanotransduction. Ann Am Thorac Soc 12 Suppl 1, S24–S29 (2015).

21. Wang, N., Butler, J. P. & Ingber, D. E. Mechanotransduction Across the Cell Surface and Through the Cytoskeleton. Science (1979) 260, 1124–1127 (1993).

22. Rosmark, O. et al. Alveolar epithelial cells are competent producers of interstitial extracellular matrix with disease relevant plasticity in a human in vitro 3D model. Sci Rep 13, (2023).

23. Huang, X. et al. Matrix Stiffness-Induced Myofibroblast Differentiation Is Mediated by Intrinsic Mechanotransduction. doi:10.1165/rcmb.2012-0050OC.

24. Bello, A. B., Kim, D., Kim, D., Park, H. & Lee, S. H. Engineering and Functionalization of Gelatin Biomaterials: From Cell Culture to Medical Applications. https://home.liebertpub.com/teb 26, 164–180 (2020).

25. Robinson, M., Douglas, S. & Willerth, S. M. Mechanically stable fibrin scaffolds promote viability and induce neurite outgrowth in neural aggregates derived from human induced pluripotent stem cells. Scientific Reports 2017 7:1 7, 1–9 (2017).

26. Eyrich, D. et al. Long-term stable fibrin gels for cartilage engineering. Biomaterials 28, 55–65 (2007).

27. Lou, J., Stowers, R., Nam, S., Xia, Y. & Chaudhuri, O. Stress relaxing hyaluronic acid-collagen hydrogels promote cell spreading, fiber remodeling, and focal adhesion formation in 3D cell culture. Biomaterials 154, 213–222 (2018).

28. Caliari, S. R. & Burdick, J. A. A practical guide to hydrogels for cell culture. Nat Methods 13, 405–414 (2016).

29. Locy, M. L., et al. Oxidative Cross-Linking of Fibronectin Confers Protease Resistance and Inhibits Cellular Migration. Sci. Signal vol. 13 https://www.science.org (2020).

30. Cruz, L. C. et al. Identification of tyrosine brominated extracellular matrix proteins in normal and fibrotic lung tissues. Redox Biol 71, 103102 (2024).

31. Guvendiren, M. & Burdick, J. A. Stiffening hydrogels to probe short- and long-term cellular responses to dynamic mechanics. Nat Commun 3, 792 (2012).

32. Caliari, S. R. et al. Stiffening hydrogels for investigating the dynamics of hepatic stellate cell mechanotransduction during myofibroblast activation. Scientific Reports 2016 6:1 6, 1–10 (2016).

33. Li, X. et al. Dynamic Stiffening Hydrogel with Instructive Stiffening Timing Modulates Stem Cell Fate In Vitro and Enhances Bone Remodeling In Vivo. Adv Healthc Mater 12, 2300326 (2023).

34. Barkauskas, C. E. et al. Type 2 alveolar cells are stem cells in adult lung. J Clin Invest 123, 3025–3036 (2013).

35. Kotton, D. N. & Morrisey, E. E. Lung regeneration: mechanisms, applications and emerging stem cell populations. Nature Medicine 2014 20:8 20, 822–832 (2014).

36. Parimon, T., Yao, C., Stripp, B. R., Noble, P. W. & Chen, P. Alveolar Epithelial Type II Cells as Drivers of Lung Fibrosis in Idiopathic Pulmonary Fibrosis. Int J Mol Sci 21, (2020).

37. Winters, N. I., Burman, A., Kropski, J. A. & Blackwell, T. S. Epithelial Injury and Dysfunction in the Pathogenesis of Idiopathic Pulmonary Fibrosis. doi:10.1016/j.amjms.2019.01.010.

38. Chambers, R. C. & Mercer, P. F. Mechanisms of alveolar epithelial injury, repair, and fibrosis. in Annals of the American Thoracic Society vol. 12 S16–S20 (American Thoracic Society, 2015).

39. Toth, A. et al. Alveolar epithelial progenitor cells drive lung regeneration via dynamic changes in chromatin topology modulated by lineage-specific Nkx2-1 activity. bioRxiv 2022.08.30.505919 (2022) doi:10.1101/2022.08.30.505919.

40. Fancy, D. A. & Kodadek, T. Chemistry for the analysis of protein-protein interactions: Rapid and efficient cross-linking triggered by long wavelength light. Proc Natl Acad Sci U S A 96, 6020–6024 (1999).

41. Bjork, J. W., Johnson, S. L. & Tranquillo, R. T. Ruthenium-catalyzed photo cross-linking of fibrin-based engineered tissue. Biomaterials 32, 2479 (2011).

42. Maina, M. B., Al-Hilaly, Y. K. & Serpell, L. C. Dityrosine cross-linking and its potential roles in Alzheimer’s disease. Front Neurosci 17, 1132670 (2023).

43. Mooney, R., Tawil, B. & Mahoney, M. Specific Fibrinogen and Thrombin Concentrations Promote Neuronal Rather Than Glial Growth When Primary Neural Cells Are Seeded Within Plasma-Derived Fibrin Gels. https://home.liebertpub.com/tea 16, 1607–1619 (2010).

44. O’Brien, F. J., Harley, B. A., Yannas, I. V. & Gibson, L. J. The effect of pore size on cell adhesion in collagen-GAG scaffolds. Biomaterials 26, 433–441 (2005).

45. Song, X. et al. A Novel Human-Like Collagen Hydrogel Scaffold with Porous Structure and Sponge-Like Properties. Polymers (Basel) 9, 638 (2017).

46. Thangprasert, A., Tansakul, C., Thuaksubun, N. & Meesane, J. Mimicked hybrid hydrogel based on gelatin/PVA for tissue engineering in subchondral bone interface for osteoarthritis surgery. Mater Des 183, 108113 (2019).

47. Liu, C., Hua, J., Ng, P. F. & Fei, B. Photochemistry of bioinspired dityrosine crosslinking. J Mater Sci Technol 63, 182–191 (2021).

48. Marquez, L. A. & Dunford, H. B. Kinetics of Oxidation of Tyrosine and Dityrosine by Myeloperoxidase Compounds I and II: IMPLICATIONS FOR LIPOPROTEIN PEROXIDATION STUDIES. Journal of Biological Chemistry 270, 30434–30440 (1995).

49. Hafidz, R. N. R. M., Yaakob, C. M., Amin, I. & Noorfaizan, A. Chemical and functional properties of bovine and porcine skin gelatin. Int Food Res J 18, (2011).

50. Loebel, C., Broguiere, N., Alini, M., Zenobi-Wong, M. & Eglin, D. Microfabrication of photo-cross-linked hyaluronan hydrogels by single- and two-photon tyramine oxidation. Biomacromolecules 16, 2624–2630 (2015).

51. Nizamoglu, M. et al. An in vitro model of fibrosis using crosslinked native extracellular matrix-derived hydrogels to modulate biomechanics without changing composition. Acta Biomater 147, 50–62 (2022).

52. Akram, K. M. et al. Live imaging of alveologenesis in precision-cut lung slices reveals dynamic epithelial cell behaviour. Nat Commun 10, 1178 (2019).

53. Sanderson, M. J. Exploring lung physiology in health and disease with lung slices. Pulm Pharmacol Ther 24, 452–465 (2011).

54. Koziol-White, C., Gebski, E., Cao, G. & Panettieri, R. A. Precision cut lung slices: an integrated ex vivo model for studying lung physiology, pharmacology, disease pathogenesis and drug discovery. Respir Res 25, 231 (2024).

55. Matera, D. L. et al. Microengineered 3D pulmonary interstitial mimetics highlight a critical role for matrix degradation in myofibroblast differentiation. Sci Adv 6, (2020).

56. Bryson, K. J. et al. Precision cut lung slices: A novel versatile tool to examine host-pathogen interaction in the chicken lung. Vet Res 51, 1–16 (2020).

57. Hesse, C. et al. Nintedanib modulates type III collagen turnover in viable precision-cut lung slices from bleomycin-treated rats and patients with pulmonary fibrosis. Respir Res 23, (2022).

58. Preuß, E. B. et al. The Challenge of Long-Term Cultivation of Human Precision-Cut Lung Slices. Am J Pathol 192, 239–253 (2022).

59. Pieretti, A. C., Ahmed, A. M., Roberts, J. D. & Kelleher, C. M. A Novel In Vitro Model to Study Alveologenesis. Am J Respir Cell Mol Biol 50, 459–469 (2014).

60. Hoffman, E. T. et al. Human alveolar hydrogels promote morphological and transcriptional differentiation in iPSC-derived alveolar type 2 epithelial cells. Sci Rep 13, 12057 (2023).

61. Chioccioli, M. et al. Stem cell migration drives lung repair in living mice. Dev Cell 59, 830–840.e4 (2024).

62. Barkauskas, C. E. et al. Type 2 alveolar cells are stem cells in adult lung. Journal of Clinical Investigation 123, 3025–3036 (2013).

63. Dupont, S. et al. Role of YAP/TAZ in mechanotransduction. Nature 474, 179–184 (2011).

64. LaCanna, R. et al. Yap/Taz regulate alveolar regeneration and resolution of lung inflammation. Journal of Clinical Investigation 129, 2107–2122 (2019).

65. Yao, C. et al. Senescence of Alveolar Type 2 Cells Drives Progressive Pulmonary Fibrosis. Am J Respir Crit Care Med 203, 707–717 (2021).

66. Wang, F. et al. Regulation of epithelial transitional states in murine and human pulmonary fibrosis. J Clin Invest 133, (2023).

67. Liang, J. et al. Reciprocal interactions between alveolar progenitor dysfunction and aging promote lung fibrosis. Elife 12, 85415 (2023).

68. Yang, H., Xu, Z., Peng, Y., Wang, J. & Xiang, Y. Integrin β4 as a Potential Diagnostic and Therapeutic Tumor Marker. Biomolecules 11, 11 (2021).

69. Blokland, K. E. C., Pouwels, S. D., Schuliga, M., Knight, D. A. & Burgess, J. K. Regulation of cellular senescence by extracellular matrix during chronic fibrotic diseases. Clin Sci (Lond) 134, 2681 (2020).

70. Hamill, K. J., Kligys, K., Hopkinson, S. B. & Jones, J. C. R. Laminin deposition in the extracellular matrix: a complex picture emerges. J Cell Sci 122, 4409 (2009).

71. Schuger, L. Laminins in lung development. Exp Lung Res 23, 119–129 (1997).

72. DeBiase, P. J. et al. Laminin-311 (laminin-6) fiber assembly by type I-like alveolar cells. Journal of Histochemistry and Cytochemistry 54, 665–672 (2006).

73. Dunsmore, S. E., Lee, Y. C., Martinez-Williams, C. & Rannels, D. E. Synthesis of fibronectin and laminin by type II pulmonary epithelial cells. American Journal of Physiology-Lung Cellular and Molecular Physiology 270, L215–L223 (1996).

74. Tan, M. L. et al. Integrin-β4 regulates the dynamic changes of phenotypic characteristics in association with epithelial-mesenchymal transition (EMT) and RhoA activity in airway epithelial cells during injury and repair. Int J Biol Sci 18, 1254 (2022).

75. Upagupta, C., Shimbori, C., Alsilmi, R. & Kolb, M. Matrix abnormalities in pulmonary fibrosis. European Respiratory Review 27, 180033 (2018).

76. Krammer, A., Craig, D., Thomas, W. E., Schulten, K. & Vogel, V. A structural model for force regulated integrin binding to fibronectin’s RGD-synergy site. Matrix Biology 21, 139– 147 (2002).

77. Smith, M. L. et al. Force-Induced Unfolding of Fibronectin in the Extracellular Matrix of Living Cells. PLoS Biol 5, e268 (2007).

78. Humphries, J. D., Byron, A. & Humphries, M. J. Integrin ligands at a glance. J Cell Sci 119, 3901–3903 (2006).

79. Takahashi, S. et al. The RGD motif in fibronectin is essential for development but dispensable for fibril assembly. J Cell Biol 178, 167 (2007).

80. Plosa, E. J., et al. β1 Integrin regulates adult lung alveolar epithelial cell inflammation. JCI Insight 5, (2020).

81. Sucre, J. M. S., et al. Alveolar repair following LPS-induced injury requires cell-ECM interactions. JCI Insight 8, (2023).

82. Wu, H. et al. Progressive Pulmonary Fibrosis Is Caused by Elevated Mechanical Tension on Alveolar Stem Cells. Cell 180, 107–121.e17 (2020).

83. Bian, F. et al. Lung endothelial cells regulate pulmonary fibrosis through FOXF1/R-Ras signaling. Nature Communications 2023 14:1 14, 1–20 (2023).

84. Zhao, W. et al. Endothelial cell-derived MMP19 promotes pulmonary fibrosis by inducing E(nd)MT and monocyte infiltration. Cell Communication and Signaling 21, 1–17 (2023).

85. Simões, F. C. et al. Macrophages directly contribute collagen to scar formation during zebrafish heart regeneration and mouse heart repair. Nature Communications 2020 11:1 11, 1–17 (2020).

86. Mondoñedo, J. R. et al. A High-Throughput System for Cyclic Stretching of Precision-Cut Lung Slices During Acute Cigarette Smoke Extract Exposure. Front Physiol 11, 530295 (2020).

87. Davidovich, N., Huang, J. & Margulies, S. S. Bioengineering the Lung: Molecules, Materials, Matrix, Morphology, and Mechanics: Reproducible uniform equibiaxial stretch of precision-cut lung slices. Am J Physiol Lung Cell Mol Physiol 304, L210 (2013).

88. Bailey, K. E. et al. Embedding of precision-cut lung slices in engineered hydrogel biomaterials supports extended ex vivo culture. Am J Respir Cell Mol Biol 62, 14–22 (2020).

89. PIV analyser. https://imagej.net/plugins/piv-analyser.

90. Han, B. et al. AFM-Nanomechanical Test: An Interdisciplinary Tool That Links the Understanding of Cartilage and Meniscus Biomechanics, Osteoarthritis Degeneration, and Tissue Engineering. ACS Biomater Sci Eng 3, 2033–2049 (2017).

91. Kawamoto, T. & Kawamoto, K. Preparation of Thin Frozen Sections from Nonfixed and Undecalcified Hard Tissues Using Kawamoto’s Film Method (2020). Methods Mol Biol 2230, 259–281 (2021).

92. Kwok, B. et al. Rapid specialization and stiffening of the primitive matrix in developing articular cartilage and meniscus. Acta Biomater 168, 235–251 (2023).

93. Al-Mayah, A., Moseley, J., Velec, M. & Brock, K. K. Sliding characteristic and material compressibility of human lung: parametric study and verification. Med Phys 36, 4625– 4633 (2009).

94. Chapman, H. A. et al. Integrin α6β4 identifies an adult distal lung epithelial population with regenerative potential in mice. J Clin Invest 121, 2855–2862 (2011).

