## Supplementary Information for "Local photo-crosslinking of native tissue matrix regulates cell function"

[1]Department of Biomedical Engineering University of Michigan, [2]Department of Materials Science and Engineering University of Michigan, [3]School of Biomedical Engineering, Science and Health Systems, Drexel University, [4]Department of Internal Medicine, University of Michigan, [5]Cellular and Molecular Biology Program, University of Michigan, *equal contribution

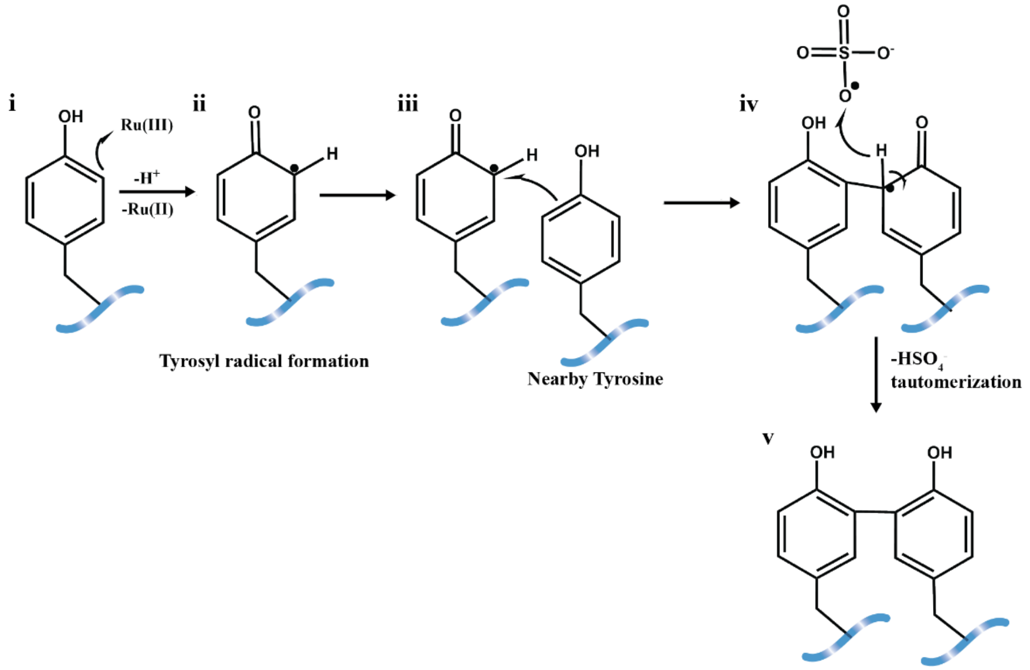


**Supplementary Figure 1**: Schematic illustrating the mechanism of dityrosine bond formation through tyrosine functional groups on most ECM proteins. Sodium persulfate (SPS) and ruthenium (Ru(II)bpy$_3^{2+}$ generate Ru(III) and sulfate radicals that oxidize tyrosines to form tyrosyl radicals (**i-ii**) that react with neighboring tyrosines (**iii**) via arene coupling. After a coupling, a hydrogen atom is lost (**iv**) to stabilize the product via tautomerization, leaving a stable dityrosine bond between two tyrosine residues (**v**).

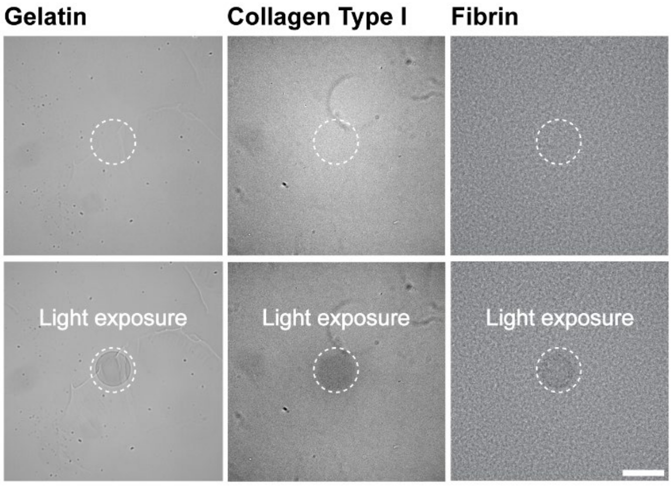


**Supplementary Figure 2:** Brightfield images of photo-crosslinking of gelatin, collagen type I, and fibrin hydrogels before and after local exposure with blue light (scale bar 100 µm).

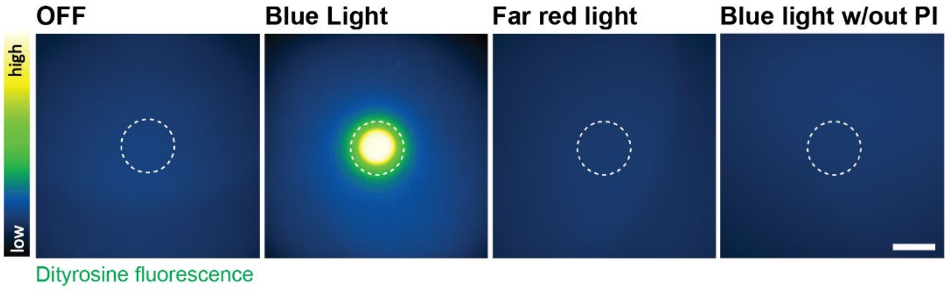


**Supplementary Figure 3:** Representative heat maps of gelatin hydrogels before (OFF) and after (ON) local exposure to blue light, far red light (670 nm) and blue light without photo-initiator (PI) (scale bar 100 µm).

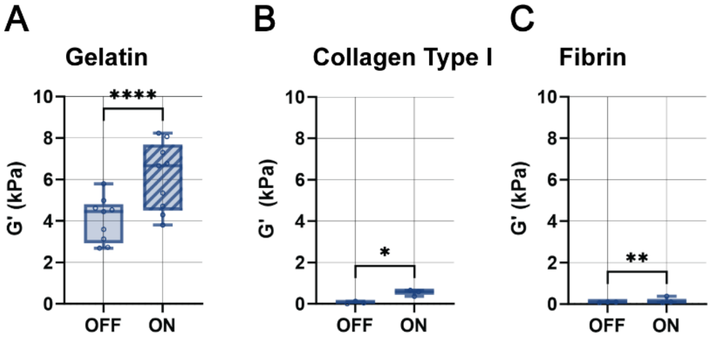


**Supplementary Figure 4:** Quantification of storage modulus (G') before (OFF) and after (ON) photo-crosslinking of **A.** gelatin , **B.** collagen type I, and **C.** fibrin hydrogels (n=9 measurements per group (gelatin), n=3 measurements per group (collagen), n=3 measurements per group (fibrin), mean ± s.d., ****$p \leq 0.0001$, **$p \leq 0.01$, *$p \leq 0.05$ by Student's two-tailed paired Student's *t*-test).

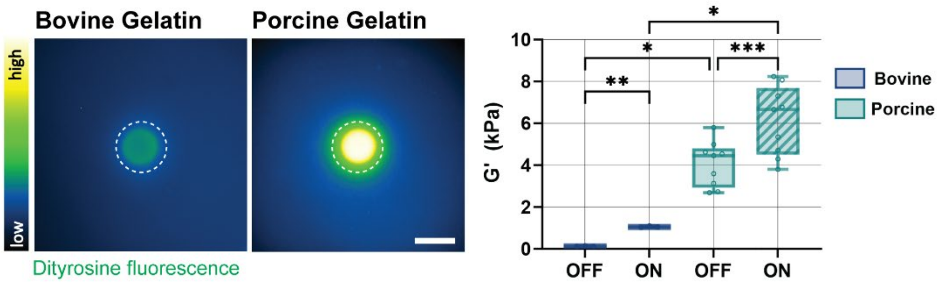


**Supplementary Figure 5:** Representative heat maps of bovine and porcine gelatin hydrogels (150mg/mL, 0.13mM Ru, ~20mW/cm$^2$ blue light for 3 minutes) and quantification of storage modulus (G') before (OFF) and after (ON) photo-crosslinking of bovine (n=3 measurements per group) and porcine (n=6 measurements per group) gelatin hydrogels, (mean ± s.d., ****$p \leq 0.0001$, ***$p \leq 0.001$, **$p \leq 0.01$, one way ANOVA with Tukey's multiple comparisons test, scale bar 100 µm).

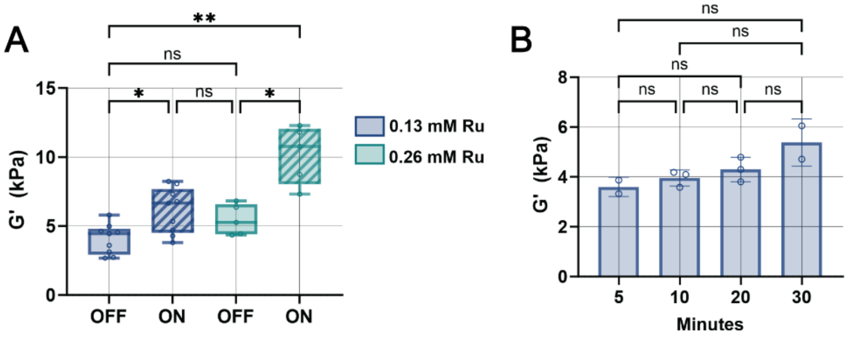


**Supplementary Figure 6: A.** Quantification of storage moduli (G') of bovine gelatin hydrogels (150 mg/mL) before (OFF) and after (ON) exposure to blue light (~20mW/cm$^2$ blue light for 3 minutes) with 0.13 mM and 0.26 mM ruthenium (Ru) (n=9 hydrogels per group (0.13mM) and n=5 hydrogels per group (0.26mM), mean ± s.d, ****p ≤ 0.0001, one-way ANOVA with Tukey's multiple comparisons test). **B.** Quantification of storage moduli (G') of gelatin hydrogels (bovine, 150 mg/mL) after incubation in PBS containing 0.13 mM Ru for 5, 10, 20 and 30 min prior exposure to blue light (n=2 hydrogels per group (5 min, 30 min) and n=3 hydrogels per group (10 min, 20 min), mean ± s.d, ****p ≤ 0.0001, one-way ANOVA with Tukey's multiple comparisons test).

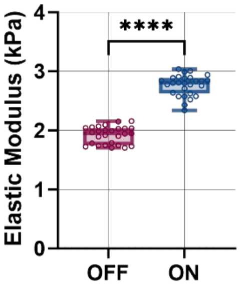


**Supplementary Figure 7:** Quantification of local Young's moduli of gelatin hydrogels (150 mg/mL) before (OFF) and after (ON) upon exposure to blue light (0.13 mM Ru, 20 mW/cm$^2$ for 3 minutes) by atomic force microscopy nanoindentation (spherical tip size of 6.79 µm, n = 27 measurements from 1 representative hydrogel per condition, mean ± s.d., ****p<0.001, two-tailed unpaired Student's *t*-test with Welch's correction).

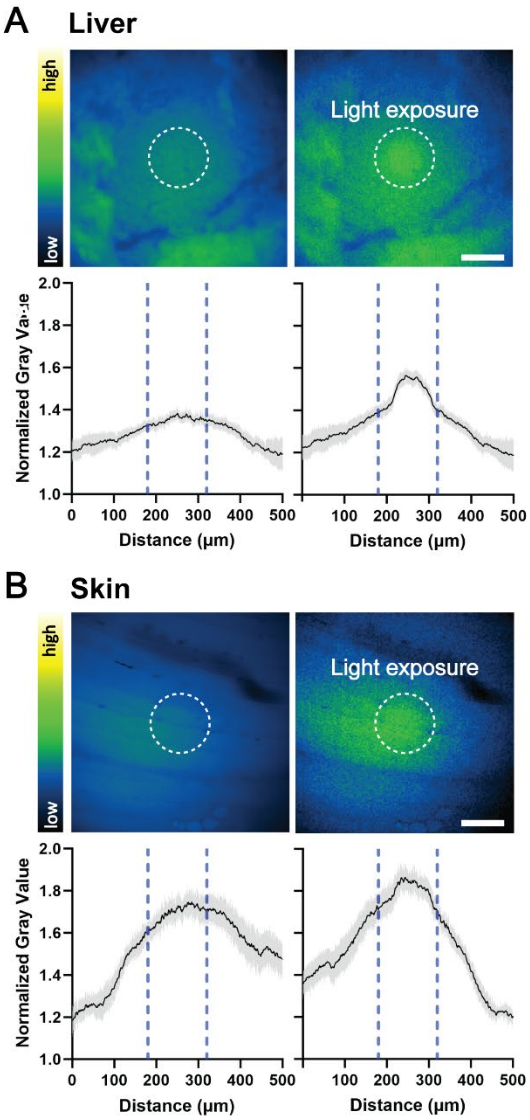


**Supplementary Figure 8:** Photo-crosslinking in other tissues. **A**. Representative heat maps and quantification of dityrosine fluorescence of murine liver tissue before (CTRL) and after (STIFF) blue light exposure (n=3 slices from one mouse, mean+SEM, 0.13 mM Ru, 20 mW/cm$^2$ for 3 minutes, scale bar 100 μm) **B.** Representative heat maps and quantification of dityrosine fluorescence of murine skin tissue before (CTRL) and after (STIFF) blue light exposure (n=3 slices from one mouse, mean+SEM, 0.13 mM Ru, 20 mW/cm$^2$ for 3 minutes, scale bar 100 μm)

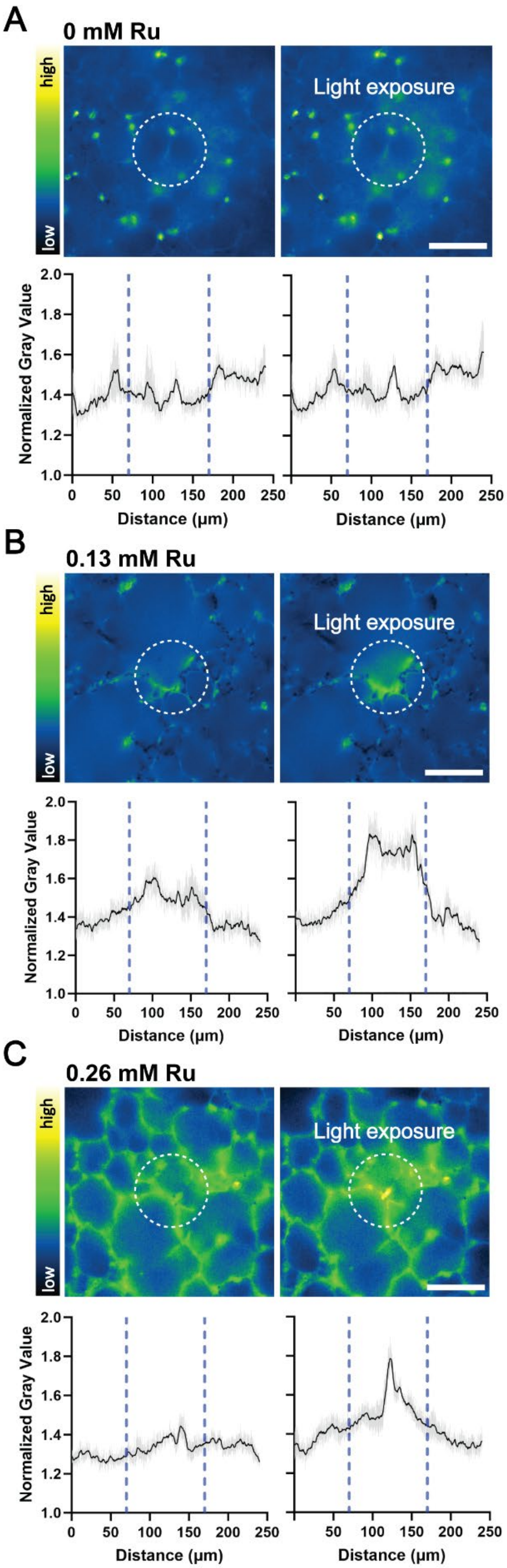
A
0 mM Ru
high
low
Light exposure
Normalized Gray Value
Distance (µm)
B
0.13 mM Ru
high
low
Light exposure
Normalized Gray Value
Distance (µm)
C
0.26 mM Ru
high
low
Light exposure
Normalized Gray Value
Distance (µm)

**Supplementary Figure 9:** Representative heat maps and quantification of dityrosine fluorescence of murine PCLS before (CTRL) and after (STIFF) blue light exposure with 0 mM, 0.13 mM, and 0.26 mM Ru) (n=1 PCLS from one mouse, mean+SEM, scale bar 100 μm).

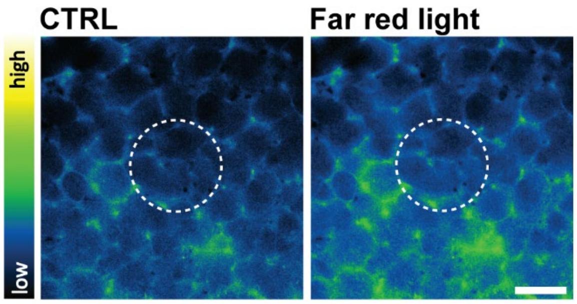


**Supplementary Figure 10:** Representative heat maps of dityrosine fluorescence before (CTRL) and upon exposure with far red light (scale bar 100 μm).

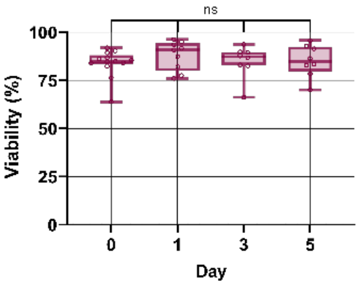


**Supplementary Figure 11:** Quantification of PCLS cell viability for up to 5 days of culture (Hoechst and Ethidium Homodimer-1, n = 13 images (day 0), 9 images (day 1), 8 images (day 3 and 5) from 3 mice, mean ± s.d., ns = no statistical difference by ANOVA with Tukey's multiple comparisons test.

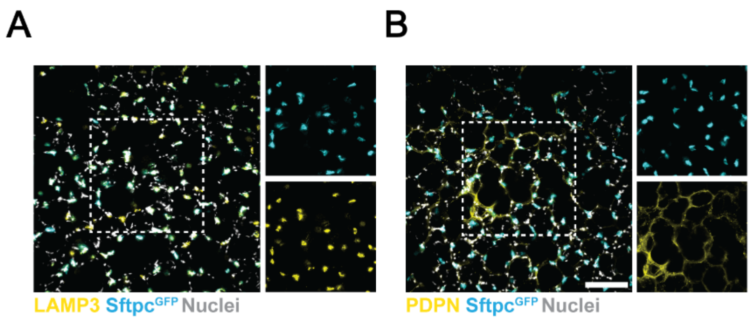


**Supplementary Figure 12: A.** Representative fluorescent images of LAMP3 staining in freshly isolated (d0) PCLS (scale bar 100 μm). **B.** Representative fluorescent images PDPN staining in freshly isolated (d0) PCLS (scale bar 100 μm).

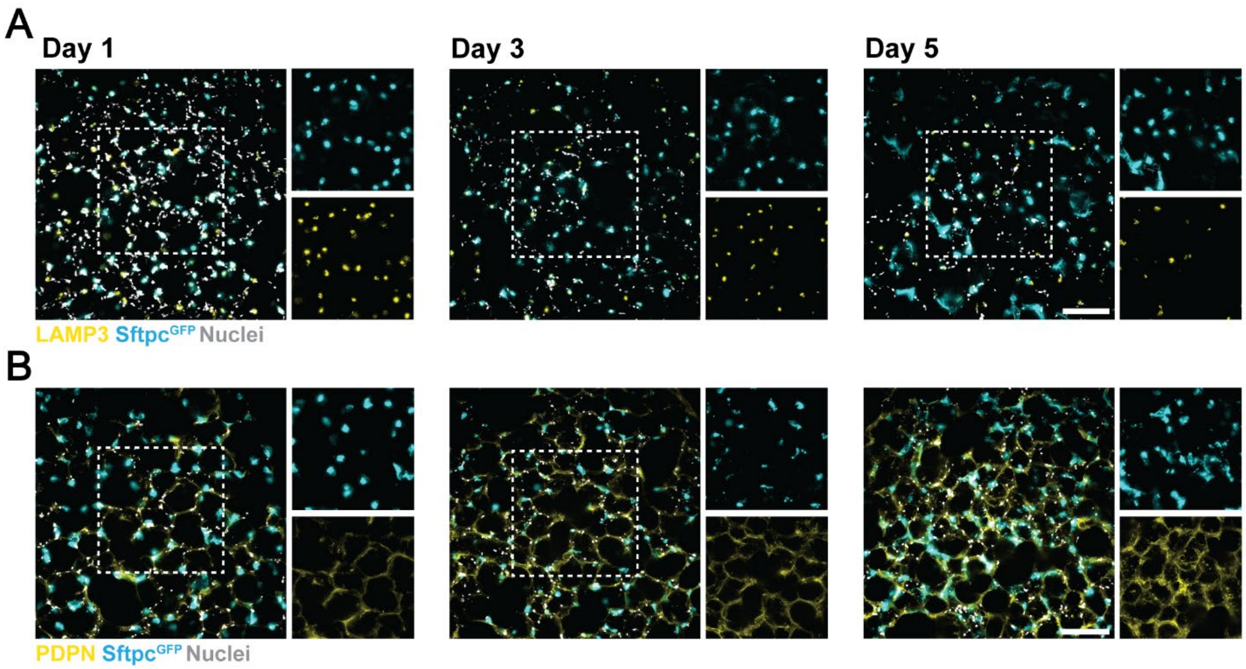


**Supplementary Figure 13:** Representative fluorescent images of **A.** LAMP3 and **B.** PDPN staining in CTRL PCLS over a 5-day culture period (scale bar 100 μm).

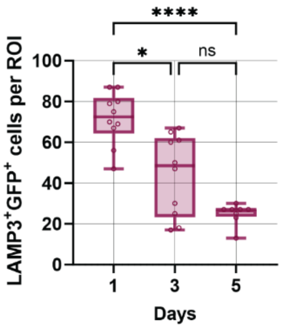


**Supplementary Figure 14:** Quantification of AT2-specific marker lysosomal associated membrane protein 3 (LAMP3) positive cells in murine PCLS up to day 5 day after PCLS preparation (n = 10 images per timepoint from 1 mouse, mean ± s.d., ****p ≤ 0.0001, *p ≤ 0.05, ns = not significant by one-way ANOVA with Tukey's multiple comparisons test).

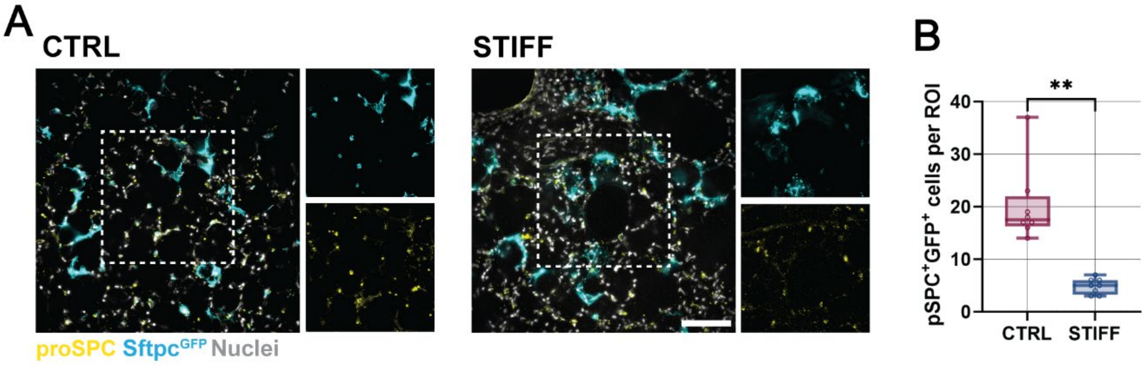


**Supplementary Figure 15: A.** Representative fluorescent images of proSPC staining in CTRL and STIFF regions in murine PCLS at day 5 (scale bar 100 µm). **B.** Quantification of proSPC$^+$/GFP$^+$ cells in CTRL and STIFF regions (n = 8 images per group (CTRL, STIFF) from one representative mouse, mean ± s.d., **p ≤ 0.01 by two-tailed unpaired Student's *t*-test with Welch's correction).

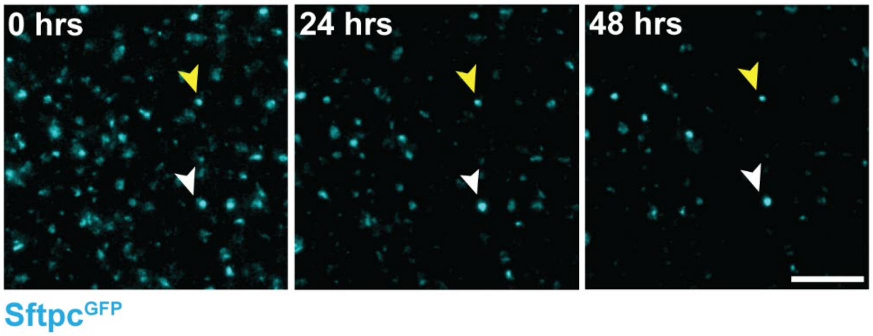


**Supplementary Figure 16:** Representative live cell fluorescent images of lineage-traced cells in CTRL regions of murine PCLS for up to 48 hours (white and yellow arrows point at two representative cells, scale bar 100 µm).

**Supplementary Video 1:** Ex vivo time-lapse video of Sftpc$^{GFP}$ cells in a CTRL and STIFF region of PCLS. Circles indicate cells over 48 hour time lapse (scale bar 100µm).

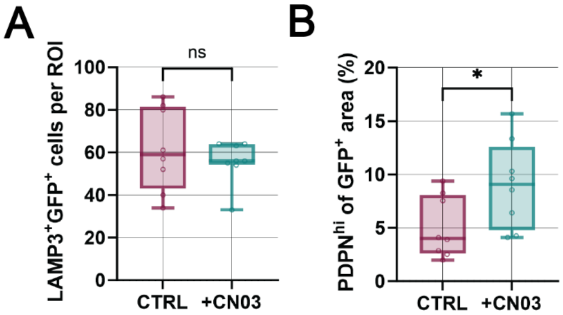


**Supplementary Figure 17: A.** Quantification of LAMP3$^{+}$GFP$^{+}$ cells expression in CTRL regions of murine PCLS without and with 5 µg/mL CN03 for 2 days (n = 8 images per group from one representative mouse, mean ± s.d, ns = not significant by unpaired *t*-test with Welch's correction). **B.** Quantification of PDPN$^{hi}$ of GFP$^{+}$ area in CTRL regions of murine PCLS without and with 5 µg/mL CN03 for 2 days (n = 8 images per group from one representative mouse (mean ± s.d, *p ≤ 0.05 by unpaired Student's *t*-test with Welch's correction).

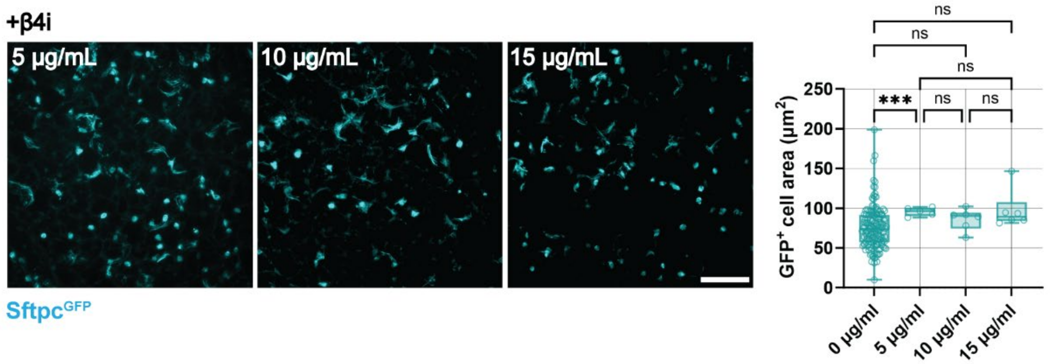


**Supplementary Figure 18:** Representative fluorescent images and quantification of GFP$^+$ cell area in STIFF regions of murine PCLS treated without (0 µg/mL) or with 5, 10 or 15 µg/mL integrin β4 function perturbing antibodies for 2 days (scale bar 100 µm, n = 149 images from 7 mice (0 µg/mL), 5 images (5 µg/mL), 6 images (10 and 15 µg/mL) from one representative mouse, mean ± s.d., ****p* ≤ 0.001, ns = not significant by one way ANOVA with Tukey's multiple comparisons test).

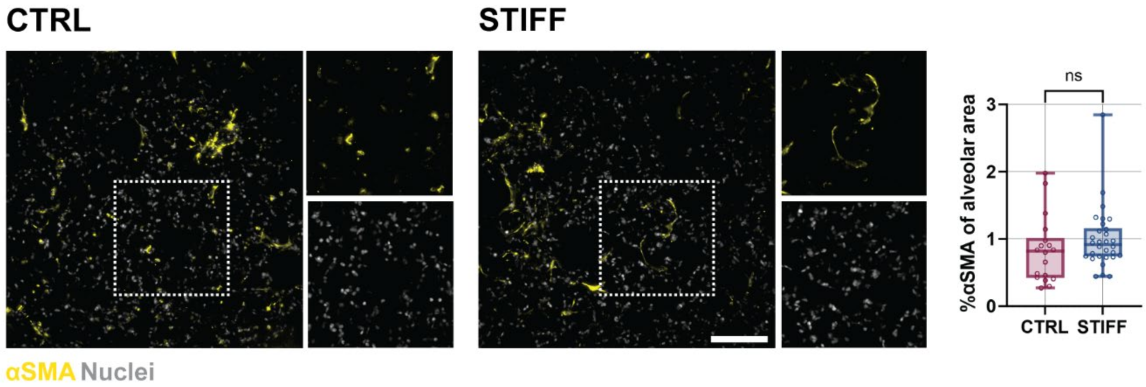


**Supplementary Figure 19:** Representative fluorescent images of alpha smooth muscle actin (αSMA) immunostaining and quantification of %αSMA per alveolar area at day 5 (n = 18 images (CTRL) and 30 images (STIFF) from 4 mice, mean ± s.d., ns = not significant by two-tailed unpaired Students *t*-test with Welch's correction, scale bar 100 µm).

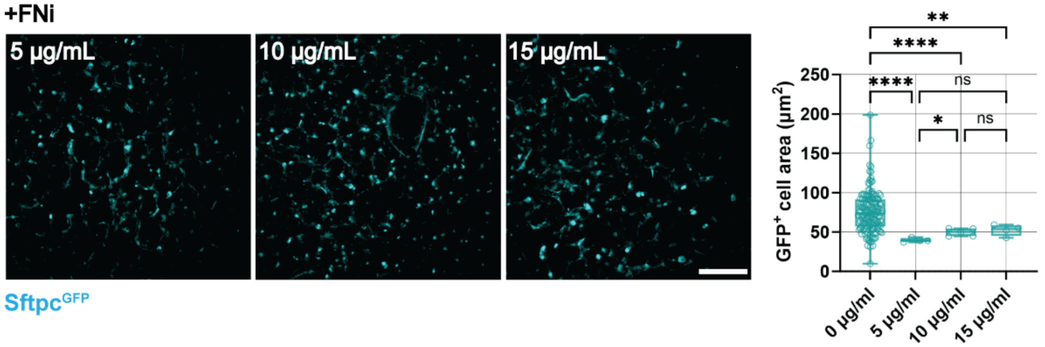


**Supplementary Figure 20:** Representative fluorescent images and quantification of GFP$^+$ cell area in STIFF regions of murine PCLS treated without (0 µg/mL) or with 5, 10 or 15 µg/mL fibronectin (FN) function perturbing antibodies for 2 days (scale bar 100 µm, n = 149 images per group from 7 mice (0 µg/mL) and n = 4-5 images from one representative mouse (5, 10 and 15 µg/mL), mean ± s.d., **p* ≤ 0.05, ***p* ≤ 0.01, *****p* ≤ 0.0001, ns = not significant by one way ANOVA with Tukey's multiple comparisons test).

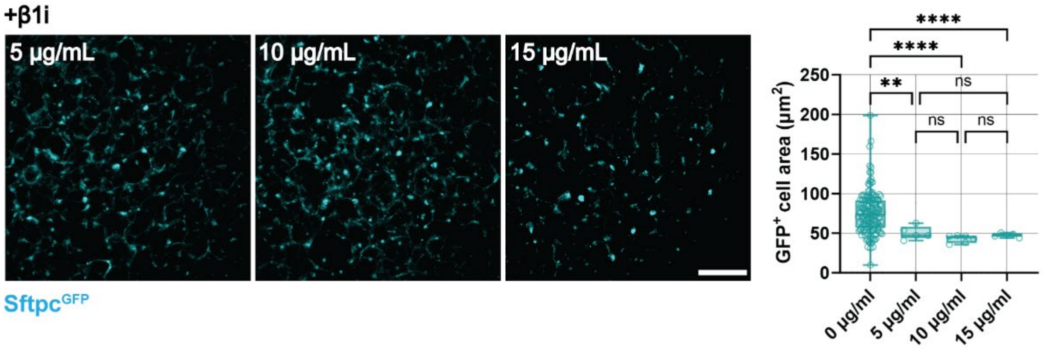


**Supplementary Figure 21:** Representative fluorescent images and quantification of GFP$^+$ cell area in STIFF regions of murine PCLS treated without (0 µg/mL) or with 5, 10 or 15 µg/mL integrin β1 function perturbing antibodies for 2 days (scale bar 100 µm, n = 149 images per group from 7 mice (0 µg/mL) and n = 5 images from one representative mouse (5, 10 and 15 µg/mL), mean ± s.d., ***p* ≤ 0.01, *****p* ≤ 0.0001, ns = not significant by one way ANOVA with Tukey's multiple comparisons test).
